# Conservation and Divergence of Oogenesis in Annual and Non-annual African Killifishes: An Oocyte Staging for the Fastest-Aging Vertebrate, *Nothobranchius furzeri*

**DOI:** 10.64898/2026.11.02.763684

**Authors:** Aarón Torres-Martínez, Essaikiammal Sodalai Muthu Konar, Martin Pšenička, Roman Franěk

## Abstract

Annual killifishes have evolved specialized reproductive strategies that allow their embryos to survive the seasonal desiccation of ephemeral habitats, yet little is known about how these adaptations arise during oogenesis. We hypothesized that traits contributing to embryonic desiccation resistance may originate from modifications in the follicular wall and egg envelope during oocyte development. Here, we provide a comparative morphological and morphometric characterization of oogenesis in the annual African turquoise killifish *Nothobranchius furzeri* and the non-annual lyretail killifish *Aphyosemion australe*, from early oocyte development through Primary Growth (PG), Secondary Growth (SG), Maturation, and Ovulation. Despite their contrasting life histories, both species exhibited a highly conserved sequence of oocyte development. Cortical alveoli accumulated before oil droplets during PG, whereas SG was characterized by the progressive coalescence of yolk globules into a homogeneous fluid yolk mass and the peripheral displacement of oil droplets and cortical alveoli. Oocyte diameter did not differ significantly between species at any developmental stage, and the follicular epithelium followed a similar developmental pattern, suggesting that its basic morphological organization is conserved despite contrasting reproductive ecologies. In contrast, greater interspecific divergence was observed in the zona pellucida, which became significantly thicker in *N. furzeri* from early SG through Maturation and exhibited distinct patterns of chorionic ornamentation between species. These findings reveal a largely conserved program of oogenesis and follicular epithelial differentiation in annual and non-annual African killifishes, while highlighting greater morphological divergence in the egg envelope. This suggests that different components of the ovarian follicle may exhibit distinct degrees of evolutionary plasticity, with modifications of the zona pellucida potentially contributing to embryo-environment interactions and desiccation resistance in annual species.

**Graphical Abstract:** 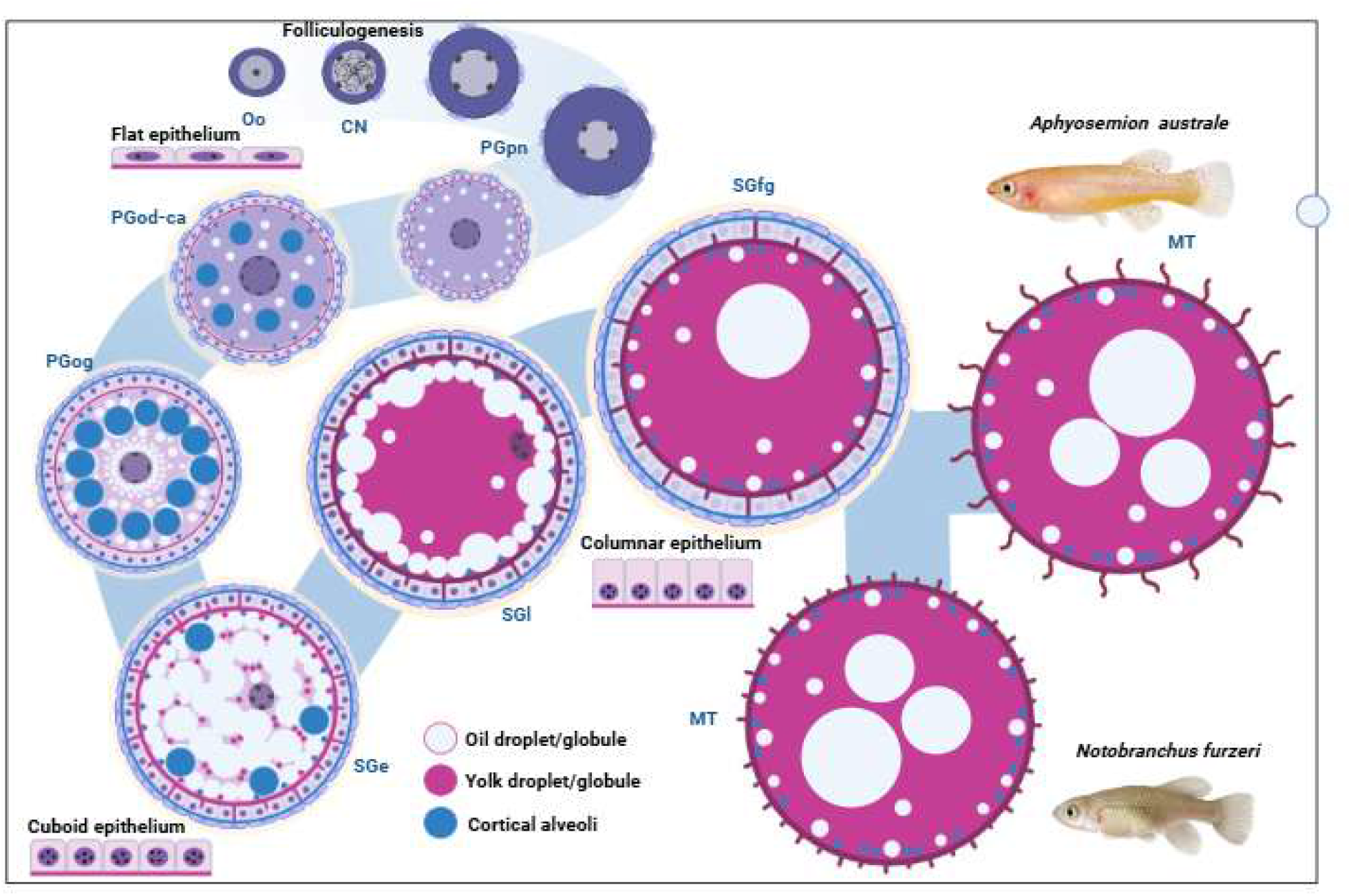

**Highlights:** Oogenesis is remarkably conserved in two contrasting killifish life histories.

Oocyte growth and cytoplasmic organization follow a shared developmental pattern

Follicular epithelial differentiation is highly conserved between species.

Conserved follicular morphology may reflect an underlying developmental constraint.

Zona pellucida thickness and ornamentation show greater evolutionary divergence.

We hypothesize that egg-envelope traits are more evolutionarily labile than follicular development.

## 1. INTRODUCTION

Oogenesis is the process through which female germ cells develop into mature oocytes capable of fertilization and supporting embryonic development. In teleost fish, this process involves a highly coordinated sequence of nuclear, cytoplasmic, and follicular changes, including oogonial proliferation and folliculogenesis, primary oocyte growth, accumulation of cortical alveoli and lipids, vitellogenesis, and final oocyte maturation (Wallace & Selman, 1981; 1990; Tyler & Sumpter, 1996; Lubzens et al., 2010). Although this general sequence is broadly conserved, teleosts display remarkable diversity in the timing and extent of these processes and in the morphology of the resulting oocytes (Grier et al., 2009; Uribe et al., 2020). Such variation is particularly evident in the accumulation and organization of cytoplasmic inclusions such as yolk, oil droplets, and cortical alveoli, as well as in the development of the follicular epithelium, theca, and egg envelope (Grier et al., 2009; Uribe et al., 2011, 2012, 2016; Torres-Martínez et al., 2017; Uribe et al., 2020). These differences may have important functional consequences, as yolk and lipid reserves determine the amount and form of maternally supplied energy available during embryogenesis, whereas cortical alveoli and the egg envelope contribute to fertilization, egg activation, and protection of the developing embryo (Wiegand et al., 1996; Lubzens et al., 2017; Perez-Atehortua et al., 2023). Variation in these oocyte components may therefore reflect adaptations to species-specific reproductive strategies and the environmental conditions experienced by embryos during development. Consequently, comparative analysis of oogenesis provides a framework for identifying both conserved mechanisms of oocyte development and lineage- or ecology-specific reproductive traits.

Aplocheilid killifishes occupy habitats ranging from relatively permanent freshwater environments to highly seasonal pools that disappear completely during the dry season, making them a particularly informative system in which to examine reproductive and developmental adaptations (Furness 2016; Reichard, 2016; Naumann & Englert, 2018). Within this clade, annualism has evolved repeatedly in association with the colonization of ephemeral aquatic habitats and is characterized by a life cycle in which adults complete reproduction during the aquatic phase, whereas embryos survive the dry season within the substrate, frequently through one or more developmental diapauses (Furness et al., 2015; Furness, 2016; Thompson et al., 2024). Consequently, the majority of developmental studies in killifishes have focused on diapause (Martin & Podrabsky, 2017; Podrabsky, & Arezo, 2017). However, much less is known about whether adaptations associated with annual life histories are already reflected during oogenesis. Although structures such as the egg envelope and perivitelline space are common features of teleost eggs, modifications in their thickness, organization, or complexity have been reported in annual killifishes and may contribute to embryo survival under prolonged terrestrial incubation (Domínguez-Castanedo et al., 2016; Papa et al., 2019). Because these structures are established from components produced or deposited during oocyte and follicular development such specializations may require corresponding modifications during oogenesis. (Domínguez-Castanedo et al., 2017; 2018).

Despite broadly conserved features of fish oocytes, including overall egg morphology and zona pellucida organization, considerable structural diversity exists among teleosts, reflecting adaptations to different reproductive strategies and environmental conditions (Honorato-Sampaio et al., 2015; Litscher et al., 2018; Hernández-Gómez et al., 2019; Mokhtar, 2024). One fundamental distinction is between pelagic eggs, which remain suspended in the water column, and demersal eggs, which are deposited on or near a substrate (Grier et al., 2018; Nande et al., 2024; Zhao et al., 2025). These strategies are associated with differences in egg envelope structure and surface specializations that affect buoyancy, substrate attachment, protection from mechanical stress, and embryo survival (Riehl et al., 1998; Sano et al., 2017; Cao et al., 2023). Yet the specific ways oocyte structures are modified in relation to life-history strategy remain incompletely characterized, particularly in annual killifishes whose embryos survive seasonal drying and prolonged desiccation (Thompson et al., 2017; Zhao et al., 2025). This knowledge gap is particularly relevant in annual killifishes, whose embryos must withstand prolonged periods of environmental desiccation. Because several structures established during oogenesis subsequently contribute directly to egg and embryo performance and survival (Sano et al., 2022; Domínguez-Castanedo, 2024), comparative analysis of oocyte architecture may reveal morphological adaptations associated with embryonic desiccation resistance. For instance, cortical alveoli participate in the cortical reaction and modification of the egg envelope following fertilization, while yolk and lipid reserves provide the resources required for embryonic development (Domínguez-Castanedo et al., 2016; 2018; 2024; Lubzens et al., 2010; 2017). Similarly, the zona pellucida (ZP) and its associated surface ornamentations form a highly variable interface between the embryo and its environment. Among killifishes, egg envelopes vary markedly in thickness, surface architecture, filaments, ridges, and other projections (Loureiro & de Sá, 1996; Arezo et al., 2005; Litscher & Thompson et al., 2017). These structures have been proposed to contribute to attachment, protection, gas exchange, and resistance to environmental stress, although their functional and evolutionary significance remains incompletely understood (Litscher & Wassarman, 2018; Papa et al., 2019). Importantly, annualism evolved repeatedly in killifishes, and chorionic characters show substantial homoplasy, making it difficult to distinguish traits associated with common ancestry from those that evolved independently under similar environmental pressures (Furness, 2016; Thompson et al., 2024). Despite the ecological and evolutionary importance of these traits, most comparative studies of killifish eggs have focused on the morphology of the mature or fertilized egg. Far less is known about how these structures arise and change during oogenesis, leaving the developmental origins of desiccation resistance largely uncharacterized (Papa et al., 2016; Domínguez-Castanedo et al., 2024). Detailed studies of ovarian development are available for a limited number of annual and non-annual cyprinodontiform fishes, including *Millerichthys robustus*, *Cyprinodon artifrons*, *Garmanella pulchra*, and *Melanorivulus rossoi* which collectively reveal extensive variation in the timing of cortical alveoli and lipid accumulation, follicular epithelium differentiation, and egg-envelope formation (Domínguez-Castanedo & Uribe, 2019; Cassel et al., 2023; Domínguez-Castanedo, 2024). These studies reveal considerable variation in the timing of cortical alveoli and lipid accumulation, follicular-wall differentiation, and egg-envelope formation. However, direct comparisons between closely related annual and non-annual African killifishes remain scarce, leaving unresolved whether the major cellular events of oogenesis are conserved across contrasting life-history strategies or differ in their developmental timing and progression (Borisov et al., 2023).

The African turquoise killifish *Nothobranchius furzeri* Jubb, 1971 inhabits ephemeral savannah pools (Fig. 1A) and exhibits the fastest sexual maturation recorded for a vertebrate, reaching reproductive maturity in as few as 14 days (Vrtílek et al., 2018; Leow et al., 2025; Franěk et al., 2026). This rapid maturation and exceptionally short adult lifespan are coupled with daily reproduction, allowing the population to complete its life cycle before seasonal habitat desiccation (Vrtílek et al., 2018; Tozzini et al., 2020). In contrast, *Aphyosemion. australe* (Rachow, 1921) is a non-annual African killifish associated with more permanent aquatic habitats and does not depend on an annual cycle of habitat desiccation and embryonic diapause (Leow et al., 2025; Fig. 1B). Despite these contrasting life histories, both species belong to the family Nothobranchiidae, providing a relatively close phylogenetic framework for examining reproductive developmental differences associated with contrasting life-history strategies (Leow et al., 2025).

**FIGURE 1.**
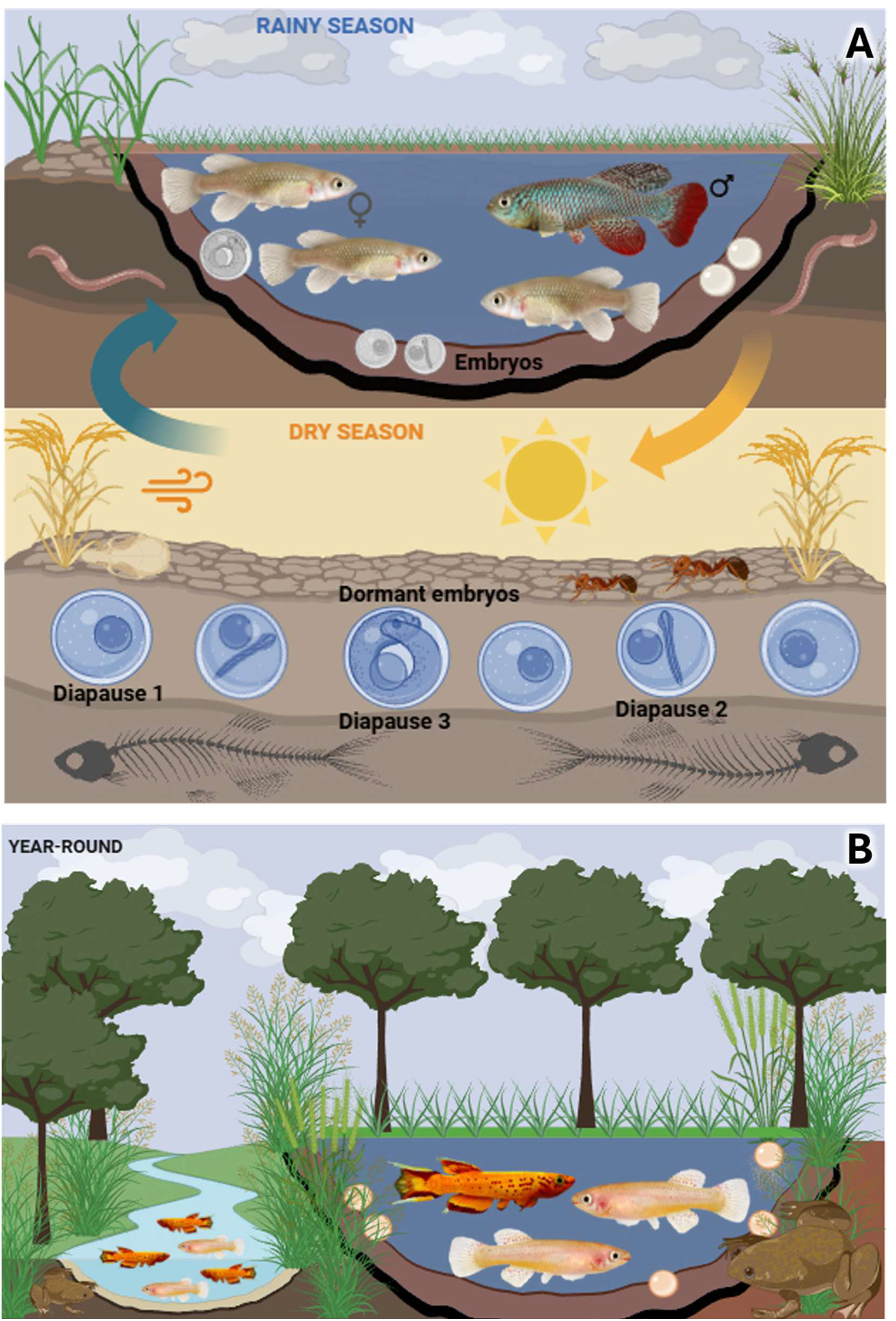
Contrasting life histories in annual killifishes and non-annual killifishes. (**A**). The annual African turquoise killifish *Nothobranchius furzeri* inhabits temporary pools that undergo seasonal drying. Adults reproduce during the rainy season by depositing eggs in the substrate. As the pools dry, the adults die, whereas the embryos survive the dry season buried in the sediment in a state of developmental arrest (diapause). Following the return of the rains, the pools refill, embryonic development resumes, and juveniles hatch, completing the annual life cycle. **B**) The non-annual African lyretail killifish *Aphyosemion australe* inhabits permanent freshwater environments, where reproduction occurs throughout the year. Embryos develop continuously under stable aquatic conditions, and successive generations coexist in the same habitat. Illustration created in BioRender by Torres-Martínez, A..

In the present study, we provide a comparative morphological and morphometric characterization of oogenesis in *N. furzeri* and *A. australe*, spanning Oogonial Proliferation and Primary Growth (PG; or previtellogenesis) through Secondary Growth (SG; or vitellogenesis), oocyte Maturation, and ovulation. Attention is given to the temporal dynamics of cortical alveolus formation, oil-droplet accumulation, and yolk deposition; changes in the germinal vesicle; differentiation of the follicular epithelium and theca; and the development, thickness, and external ornamentation of the zona pellucida. By comparing closely related annual and non-annual African killifish species, we assess the extent to which the major morphological and cellular events of oogenesis are conserved despite their contrasting life histories, while identifying differences in the timing and structural organization of oocyte and follicular development that may represent candidate features associated with annualism or, alternatively, lineage-specific traits.

## 2. MATERIALS AND METHODS

### 2.1. Fish sampling

Adult *Nothobranchius furzeri* and *Aphyosemion australe* females were obtained from our laboratory colony, originally derived from wild-caught populations in Africa maintained under controlled conditions. The species are known to inhabit permanent and ephemeral waterbodies, respectively (Fig. 1). For the turquoise killifish *N. furzeri*, strain MZCS 222 was used, which was obtained from Leibniz Institute of Aging. Jena, Germany. *A. australe* specimens were obtained from Petra-Aqua Ltd. (Prague, Czech Republic). Both species were maintained in 100 L glass aquaria kept with constant aeration and heating adjusted to 26°C. The turquoise killifish were fed with blood worms administrated three times per day until satiation. In contrast, lyretail killifish were fed dry feed (Tetra, Melle, Germany) twice daily and live food (e.g., Artemia nauplii, Ocean Nutrition) once daily, both ad libitum. For sampling, individuals were anesthetized with an overdose of tricaine methanesulfonate (MS-222; 200 mg/L, Sigma Aldrich, Burlington, United States, Cat# E10521) until loss of opercular movement and then sacrificed by decapitation. Ovaries were immediately dissected and processed for histological analyses.

### 2.2. Ethical statement and permissions

All procedures were performed in accordance with the Animal Research Committee of the Faculty of Fisheries and Protection of Waters (Vodňany, Czech Republic) and were approved by the Ministry of Agriculture of the Czech Republic (reference number: MZE-3972339723/2025-13143). Fish were maintained according to the principles based on the EU-harmonized Animal Welfare Act of the Czech Republic and principles of laboratory animal care in compliance with the national law (Act No. 246/1992 on the Protection of Animals Against Cruelty).

### 2.3. Histological preparation and light microscopy analysis

Ovarian tissue was fixed for 24 h at room temperature in Bouin’s solution (Cat# HT10132; Sigma-Aldrich, Massachusetts, US) or formaldehyde solution (Cat# 8.18708; Sigma-Aldrich). Samples were then rinsed in phosphate-buffered saline (PBS; Cat# HY-K1023; MedChemExpress, New Jersey, USA), dehydrated through a graded ethanol series (30, 50, 70, 80, and 96%), prepared from 96% or absolute ethanol supplied by Penta Chemicals Unlimited (Prague, Czech Republic), followed by absolute ethanol from the same supplier, and embedded in either JB-4 resin (Cat# EM0100; Sigma-Aldrich; Franěk et al., 2026) or Paraplast (Sigma-Aldrich; Torres-Martínez et al., 2026) for conventional histological analysis. Serial sections (5-7 µm thick) were obtained using a rotary microtome (Thermo Fisher Scientific, Waltham, MA, USA) and mounted on glass slides. Sections were stained with hematoxylin (Gill hematoxylin solution; Cat# T864.2; Carl Roth; Karlsruhe, Germany,) and eosin (Eosin Y solution; Cat# 229C.2; Carl Roth,) for general morphological characterization. Periodic acid-Schiff (PAS) staining was used to identify cortical alveoli and the zona pellucida during oocyte development, following the protocol described by Quintero-Hunter et al. (1991) using the PAS staining kit (Carl Roth, Cat# HP01.1). Mallory’s trichrome staining was also used to visualize components of the extracellular matrix and the follicles (Abcam, Cat# ab150686, Cambridge, United Kingdom). Staining solutions were prepared according to the manufacturers’ recommendations. All slides were mounted in rapid-drying mounting medium (Pertex®; Cat# 00840; HistoLab Products AB; Askim, Sweden).

Additional ovarian samples were fixed in 4% glutaraldehyde (Sigma-Aldrich, Cat# G5882) and processed for resin embedding. Briefly, samples were dehydrated in acetone, embedded in Poly/Bed 812 (Polysciences, Cat# 08791–500). Ovarian samples were cut into 2–3-mm fragments and fixed into 2.5% glutaraldehyde (Sigma-Aldrich, Cat# G5882) in PBS (MedChemExpress) and processed to obtain semithin sections (Tichopád et al., 2020; Gao et al., 2025). Ultrathin sections were cut on a Leica ultramicrotome (Leica Biosystems, Nußloch, Germany) and stained with toluidine blue. Sections, regardless of the embedding method, were examined using an Olympus IX83 microscope (Olympus, Tokyo, Japan) equipped with an Olympus DP75 camera.

### 2.4. Oocyte staging

Several systems for classifying oocyte development have been proposed in the literature. In this study, we follow the standardized and broadly applicable terminology proposed by Grier et al. (2009) and Uribe et al. (2019), in which oogenesis is organized into Stages and Steps, facilitating comparative analyses of oocyte development across fish taxa with a range of reproductive strategies.

### 2.5. Morphometric and statistical analysis

Histological micrographs were analyzed using qPath Software (Bankhead et al., 2017). Parameters measured included the oocyte diameter, the height of the follicular epithelium, and the thickness of the zona pellucida at different stages of oocyte development. For each developmental stage, at least 30 oocytes were measured across multiple ovaries (n=5). For rare stages, for which fewer oocytes were available, all suitable oocytes identified were included in the analysis. Data was analyzed with the statistical software GraphPad Prism version 10.6.1 (San Diego, United Sates). The data were not normally distributed and were therefore analyzed using the Kruskal–Wallis test followed by Dunn’s multiple-comparisons test with adjusted P values. For clarity, only biologically relevant comparisons between germ cells at the same developmental stage are shown in the figure; these comparisons are presented irrespective of statistical significancem

## 3. RESULTS

### 3.1. Oogonial proliferation (OP) stage

In *N. furzeri*, oogonial proliferation occurred predominantly in the ovarian primordium during sex differentiation (Fig. 2A-C). Dividing oogonia were observed within the germinal epithelium of early ovaries (Fig. 2D, E) but were rarely detected in adult specimens. In *A. australe*, oogonia occurred in clusters within the germinal epithelium, occasionally forming small nests. Oogonial proliferation was more frequent in juvenile ovaries (±3 months), where they were observed forming germinal nests (Fig. 2F) than in older specimens; by 6 months to 1 year of age, oogonia were sparsely distributed throughout the germinal epithelium (Fig. 2G). Oogonia measured 9.52 ± 1.47 µm in diameter in *N. furzeri* and 10.36 ± 1.64 µm in *A. australe*. In both species, oogonia possessed a single prominent nucleolus located centrally within the nucleus, with chromatin aggregates distributed along the nuclear periphery.

**FIGURE 2.**
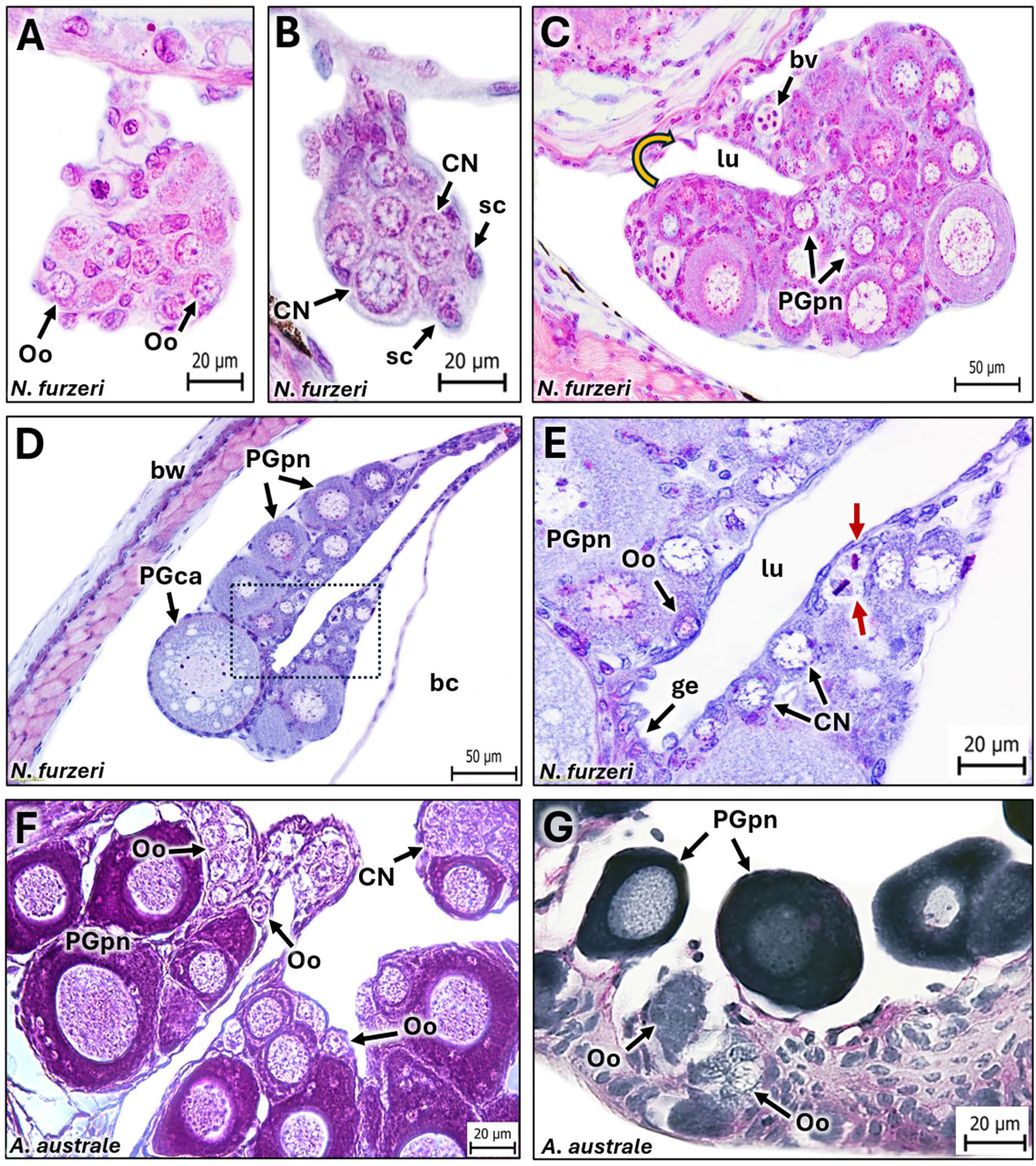
Oogonial proliferation in annual and non-annual killifishes. Ovarian differentiation and oogonial proliferation in the annual killifish *Nothobranchius furzeri* (**A-E**) and the non-annual killifish *Aphyosemion australe* (**F, G**). **(A-C)** Sequential stages of ovarian differentiation in *N. furzeri*, from the ovarian primordium to the closure of the ovarian cavity (yellow curved arrow in C). (**D, E**) Developing ovaries of juvenile *N. furzeri* showing intense oogonial proliferation within the germinal epithelium. **(E)** Higher magnification of the boxed region in (D). Mitotically dividing oogonia are indicated by red arrows. (**F, G**) Adult ovary of *A. australe*, showing widespread oogonial proliferation in the germinal epithelium. Oogonia occur either as isolated cells or as germ cell nests. **Abbreviations:** body cavity (bc), body wall (bw), Chromatin-Nucleolus oocyte (CN), germ cell nest (gn), germinal epithelium (ge), ovarian lumen (lu), oogonium (Oo), Primary-Growth perinucleolar oocyte (PGpn), and somatic cell (sc). (**A-F**) Hematoxylin and eosin staining. **(G)** Periodic acid–Schiff staining. All the sections displayed are transversal.

### 3.2. Chromatin-nucleolus (CN) stage

In both *N. furzeri* and *A. australe,* Chromatin Nucleolus (CN) oocytes were characterized by an enlarged nucleus containing chromosomes and DNA threads extending throughout the germinal vesicle (GV; or nucleus) and scarce hyaline cytoplasm. CN oocytes measured 14.20 ± 2.83 µm in *N. furzeri* and 12.09 ± 1.63 µm in *A. australe*. CN oocytes were often observed as germinal nests. Nests of CN oocytes in *N. furzeri* were observed only during the ovarian development (Fig. 2B, E), while in *A. australe* were observed in developing, juvenile, and adult ovaries (Fig. 2F, G). In both species, the cytoplasm of CN oocytes was scant, and the nucleus was markedly enlarged with conspicuous chromatin threads. Pre-follicular cells were observed at the periphery of CN oocytes; however, folliculogenesis was incomplete so no surrounding follicular layer was observed at this stage.

### 3.3. Primary Growth (PG) stage

In both *N. furzeri* and *A. australe*, PG encompassed several steps distinguishable by nuclear and cytoplasmic morphological signatures. However, not all steps described for this Stage by Grier et al. (2009) were observed in both species.

#### 3.3.1. One-nucleolus (PGon) *step*

Despite extensive screening of histological sections from multiple ovaries, we were unable to identify the one-nucleolus stage (PGon) as a discrete stage in either species. In teleosts, PGon is characterized by oocytes with a centrally located GV containing a single large nucleolus, together with the onset of cytoplasmic basophilia. However, oocytes exhibiting other morphological characteristics consistent with this stage, including oocyte diameter, cytoplasmic basophilia, and GV size, already contained multiple nucleoli distributed along the periphery of the GV.

#### 3.3.2. Multiple-nucleoli (PGmn) *step*

The subsequent multiple-nucleoli stage (PGmn) is characterized by the presence of numerous nucleoli of variable sizes within the germinal vesicle (GV), initially distributed throughout the nucleoplasm. This stage is rarely observed in fish. In *N. furzeri* and *A. australe* no oocytes displaying the typical PGmn morphology were identified; instead, oocytes within the expected size range only exhibited prominent aggregates of condensed chromatin through the GV.

#### 3.3.3. Perinucleolar (PGpn) *step*

Perinucleolar oocytes had a diameter of 38.67 ± 13.05 µm in *N. furzeri* and 32.95 ± 10.06 µm in *A. australe*. Early perinucleolar oocytes exhibited strongly basophilic cytoplasm, which gradually became more eosinophilic as oocyte growth progressed (Fig. 3A–E). Throughout this stage, oocytes displayed multiple nucleoli of variable sizes distributed along the periphery of the GV (Fig. 3B, F, E). The GV remained spherical and increased in size as the oocytes grew, while maintaining the characteristic arrangement of lampbrush chromosomes (Fig. 3B, E). In both species, the follicular epithelium consisted of a single layer of flattened follicular cells surrounding the oocyte, with a heigh of 1.702 ± 0.36 µm in *N. furzeri* and 1.21 ± 0.30 µm in *A. australe* (Fig. 3B, E).

**FIGURE 3.**
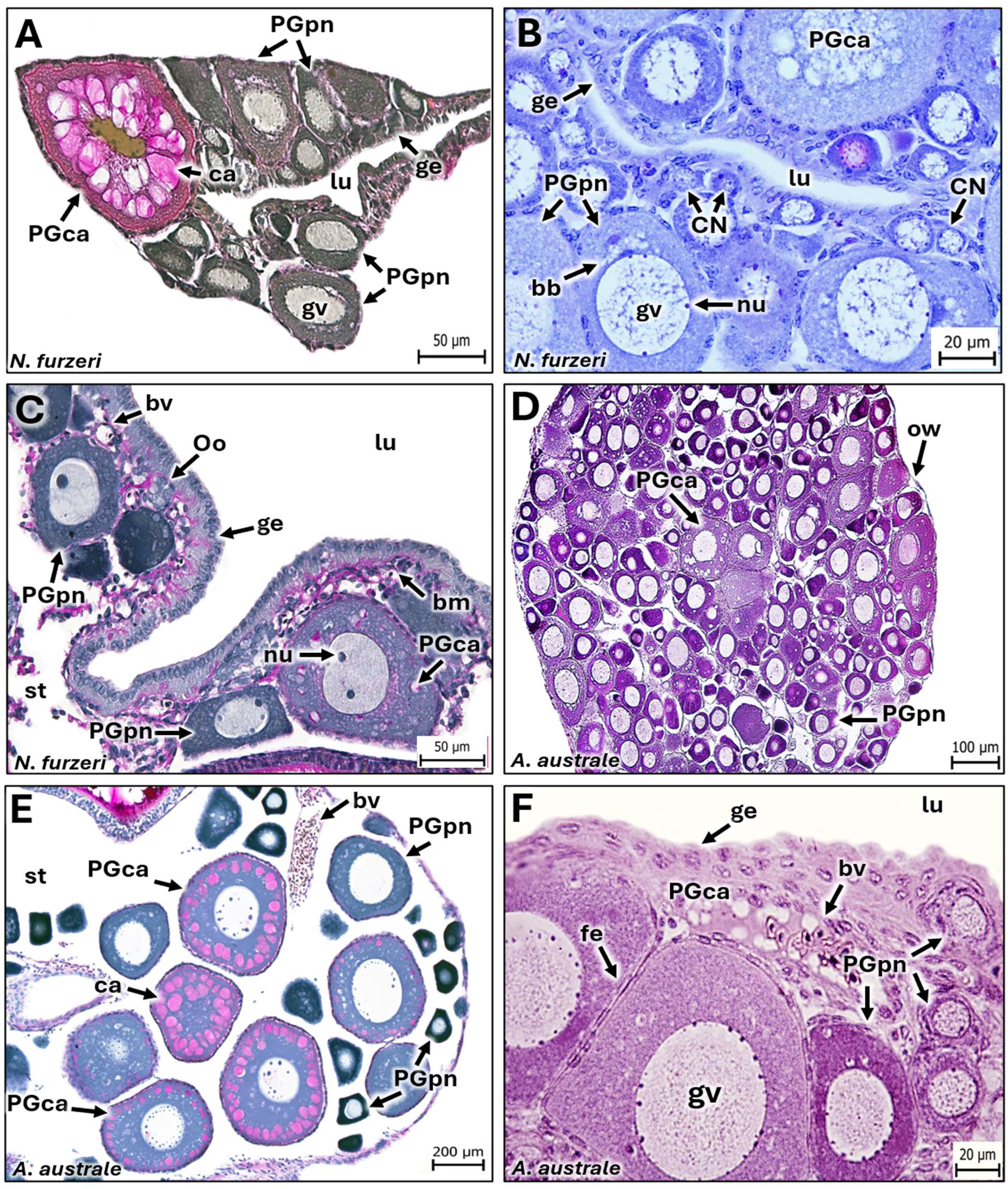
Early oocyte development in annual and non-annual killifishes: Chromatin-Nucleolus to Primary Growth cortical alveoli phases. Oocyte development in the annual killifish *Nothobranchius furzeri* (A-C) and the non-annual killifish *Aphyosemion australe* (D-F). (**A**-**F**) The earliest identifiable Primary-Growth oocytes are associated with the germinal epithelium and are characterized by a large germinal vesicle surrounded by a thin rim of cytoplasm. Distinct one-nucleolus (PGon) and multiple-nucleoli (PGmn) steps are absent, with nucleoli becoming arranged at the periphery of the germinal vesicle immediately after the chromatin-nucleolus (CN) stage. (**A**, **D**-**F**) During the Primary-Growth cortical alveoli step (PGca), PAS-positive cortical alveoli first appear in the perinuclear cytoplasm and progressively increase in size and number. PGpn and PGca oocytes are surrounded by a simple squamous follicular epithelium with an inconspicuous theca. (**A**, **B**) Developing ovaries of juvenile *N. furzeri*. (**D**) Developing ovary of a juvenile *A. australe* (approximately 1.5 months old). (**C**, **E**, **F**) Mature ovaries showing PGpn and PGca oocytes next to the ovarian lumen and germinal epithelium. **Abbreviations:** Balbiani body (bb), basement membrane (bm), blood vessel (bv), chromatin-nucleolus oocyte (CN), cortical alveoli (ca), follicular epithelium (fe), germinal epithelium (ge), germinal vesicle (gv), nucleolus (nu), oogonium (Og), ovarian lumen (lu), ovarian stroma (st), ovarian wall (ow), Primary-Growth cortical alveoli oocyte (PGca), Primary-Growth multiple-nucleoli oocyte (PGmn), and Primary-Growth perinucleolar oocyte (PGpn). (**A**, **C**, **E**) Periodic acid-Schiff and metanil yellow staining. (**B**, **D**, **F**) Hematoxylin and eosin staining.

#### 3.3.4. Cortical alveoli (PGca) step

This stage was characterized by the progressive accumulation of cortical alveoli in both species. PGca oocytes exhibited a broad size range, with a mean diameter of 74.78 ± 13.66 µm in *N. furzeri* (Fig. 4A, B) and 71.14 ± 19.60 µm in *A. australe* (Fig. 5A, C). Throughout the PGca step, the germinal vesicle retained multiple nucleoli of variable sizes distributed along its periphery (Figs. 4C, D, E; 5A–D) In *N. furzeri*, cortical alveoli first appeared as small inclusions surrounding the GV (Fig. 4A–C) and progressively increased in size and abundance during oocyte growth (Fig. 4D–F). Similarly, in *A. australe*, PAS-positive cortical alveoli first appeared as small inclusions surrounding the GV (Fig. 5A, B), preceding the appearance of the first oil droplets (Fig. 5C). As oocyte development progressed, cortical alveoli increased in size and coalesced into larger vesicles containing fibrillar material (Fig. 5D, E). In both species, in HE- stained ovaries, cortical alveoli appeared as empty vesicles, whereas in PAS-stained sections they exhibited a magenta reaction and contained PAS-positive fibrillar material. Changes in the follicular wall accompanied the accumulation of cortical alveoli. In both species, the follicular epithelium transitioned from a flattened to a cuboidal morphology during this stage (Figs. 4C–F;.5E–F) The epithelium had an average height of 2.29 ± 0.50 µm in *N. furzeri* and 1.28 ± 0.22 µm in *A. australe.* The zona pellucida first became distinguishable as a thin, homogeneous layer, with recognizable filaments extending toward the follicular epithelium. The theca became discernible only in more advanced oocytes, where it appeared as a thin layer of flattened cells.

**FIGURE 4.**
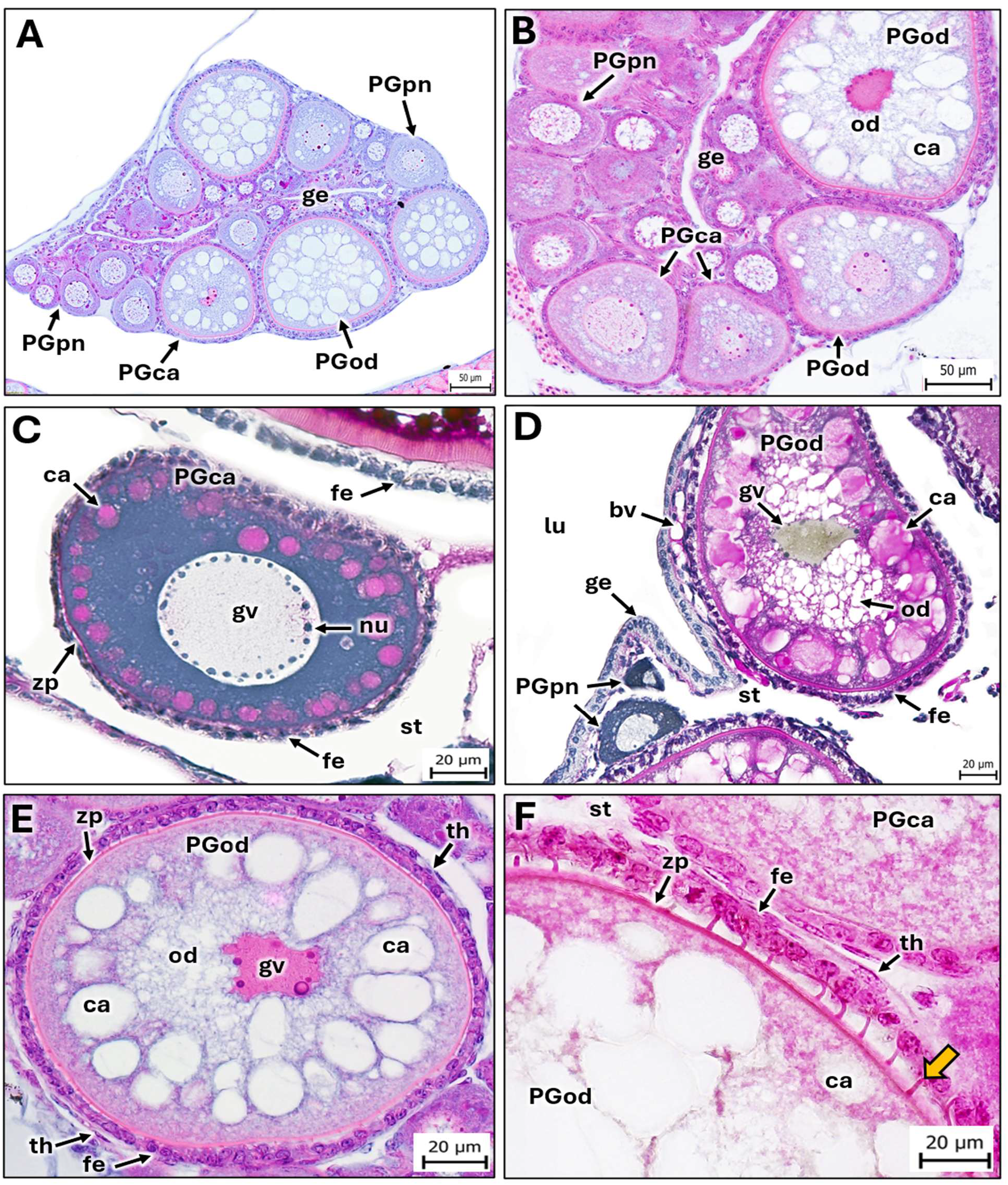
Primary Growth in the annual killifish *Nothobranchius furzeri*: cortical alveoli and oil droplet steps. Representative stages of primary oocyte growth in *N. furzeri*. (**A**-**C**). Cortical alveoli first appear in the perinuclear cytoplasm surrounding the germinal vesicle and precedes the deposition of oil droplets. (**D**). As oocyte growth progresses, oil droplets accumulate in the perinuclear region, progressively displacing the enlarging cortical alveoli toward the oocyte cortex. (**C**, **E**, **F**). The zona pellucida appears as early as the Primary Growth cortical alveoli step (PGca) but becomes more evident during the Primary-Growth oil droplet step (PGod), where it appears as a thin layer bearing fine projections extending toward the follicular epithelium (yellow arrow). Panels A and B show a developing ovary. **Abbreviations:** basal membrane (bm), blood vessel (bv), Chromatin-Nucleolus oocyte (CN), follicular epithelium (fe), germinal epithelium (ge), germinal vesicle (gv), nucleoli (nu), ovarian lumen (lu), ovarian stroma (st), cortical alveoli (ca), oil droplet (od), zona pellucida (zp), theca (th), ovarian wall (ow), Primary-Growth cortical alveoli oocyte (PGca), Primary-Growth oil droplet oocyte (PGod), and Primary-Growth perinucleolar oocyte (PGpn). (**A**, **B**, **E**, **F**) Hematoxylin and eosin staining. (**C**, **D**) Periodic acid-Schiff and metanil yellow staining.

**FIGURE 5.**
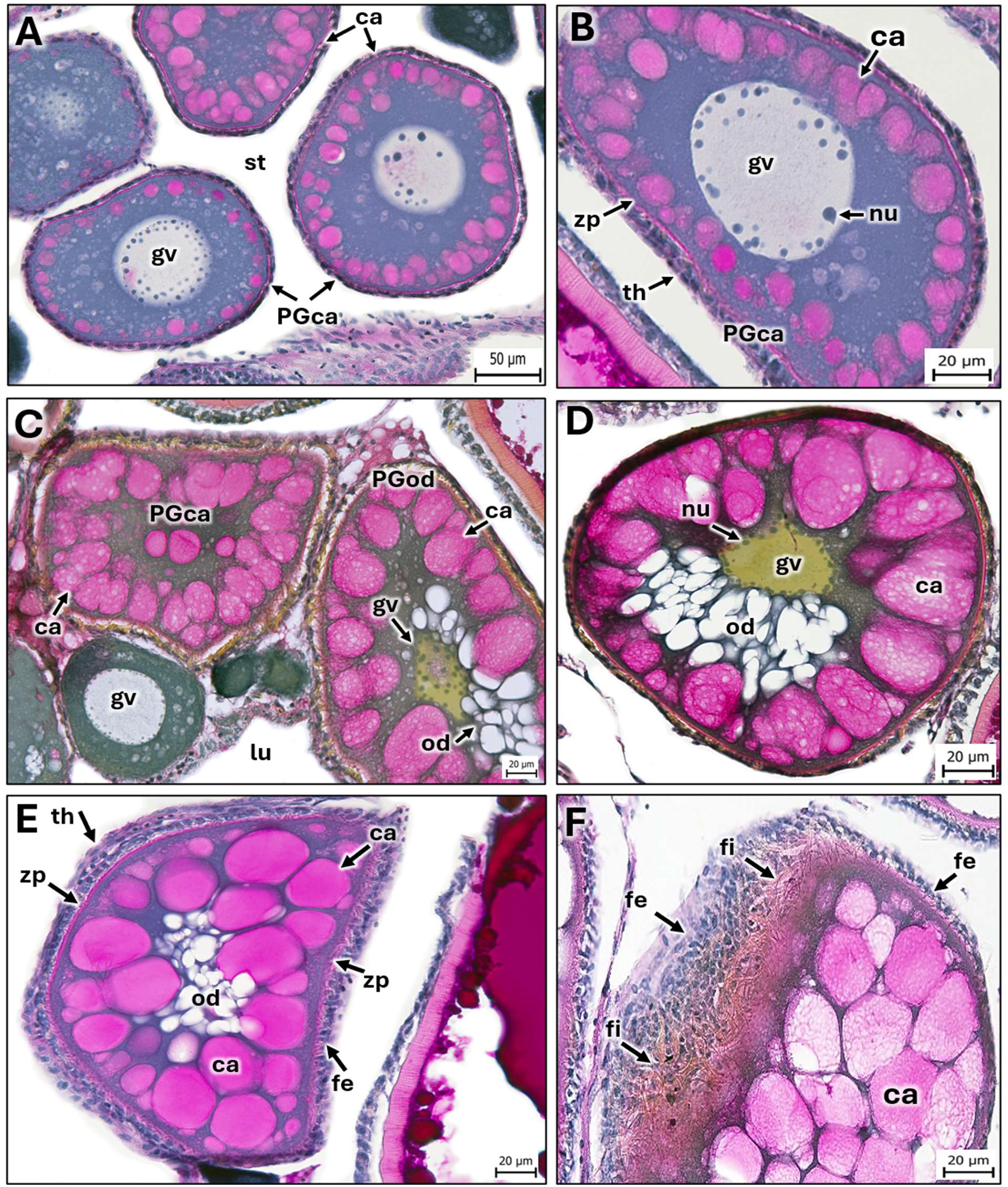
Primary Growth (PG) in the non-annual killifish *Aphyosemion australe*: cortical alveoli and oil droplet steps. (**A**, **B**) Primary-growth cortical alveoli oocytes (PGca). Cortical alveoli first appear in the perinuclear cytoplasm surrounding the germinal vesicle and precedes the deposition of oil droplets. They are Periodic acid-Schiff (PAS)-positive and, in irregularly shaped oocytes, tend to be concentrated around the middle region of the germinal vesicle. As PG progresses, cortical alveoli increase in both size and number and develop a fibrillar appearance. (**C**–**F**) The appearance of the first oil droplets marks the Primary-Growth oil droplet stage (PGod). Oil droplets arise in the perinuclear region and progressively displace the cortical alveoli toward the oocyte cortex. Consequently, the germinal vesicle becomes eccentrically positioned and adopts an irregular shape while multiple nucleoli remain conspicuous. The zona pellucida first appears as a thin PAS-positive layer and develops fine projections extending toward the follicular epithelium, which remains as a simple squamous layer throughout this stage. (**F**) Fine projections arising from the zona pellucida are particularly well illustrated. **Abbreviations:** cortical alveoli (ca), follicular epithelium (fe), germinal vesicle (gv), nucleolus (nu), oil droplet (od), ovarian lumen (lu), Primary-Growth cortical alveoli oocyte (PGca), Primary-Growth oil droplet oocyte (PGod), stroma (st), theca (th), zona pellucida (zp), and zona pellucida projections (fi). (**A–F**) Periodic acid-Schiff and metanil yellow staining.

#### 3.3.5. Oil droplet (PGod)step

These oocytes had an average diameter of 115.3 ± 34.63 µm in *N. furzeri* and 139.9 ± 43.64 µm in *A. australe*. In both species, oil droplets first appeared at the periphery of the GV and progressively increased in number and size throughout the ooplasm. As oil deposition proceeded, cortical alveoli became displaced toward the periphery of the oocyte (Figs. 4D, E; 5C–E). During this stage, the GV became eccentrically positioned and adopted an irregular shape, while the nucleoli remained distributed along its periphery (Figs. 4D, E; 5C, D). The follicular epithelium remained cuboidal throughout this stage (Figs. 4E, F; 5E). At this step, the mean height of the follicular epithelium was 5.22 ± 1.16 µm in *N. furzeri* and 3.75 ± 1.40 µm in *A. australe*. The zona pellucida became more prominent in both species, and the developing accessory filaments were first clearly distinguishable at this stage (Figs. 4F; 5F). The zona pellucida had a mean thickness of 1.18 ± 0.31 µm in *N. furzeri* and 1.70 ± 0.40 µm in *A. australe*.

### 3.4. Secondary Growth (SG) stage

SG was characterized by the progressive accumulation of yolk, from yolk granules to yolk globules that coalesced in a mass of homogeneous fluid yolk in both species.

#### 3.4.1. Early secondary growth step (SGe)

In *N. furzeri*, the onset of SG was characterized by the massive coalescence of oil droplets and the deposition of yolk granules as diffuse eosinophilic material surrounding the oil droplets (Fig. 6A–F). SGe oocytes in *N. furzeri* had an average diameter of 319.9 ± 71.12 µm, while in *A. australe* the average diameter was 263.3 ± 24.60 µm.

**FIGURE 6.**
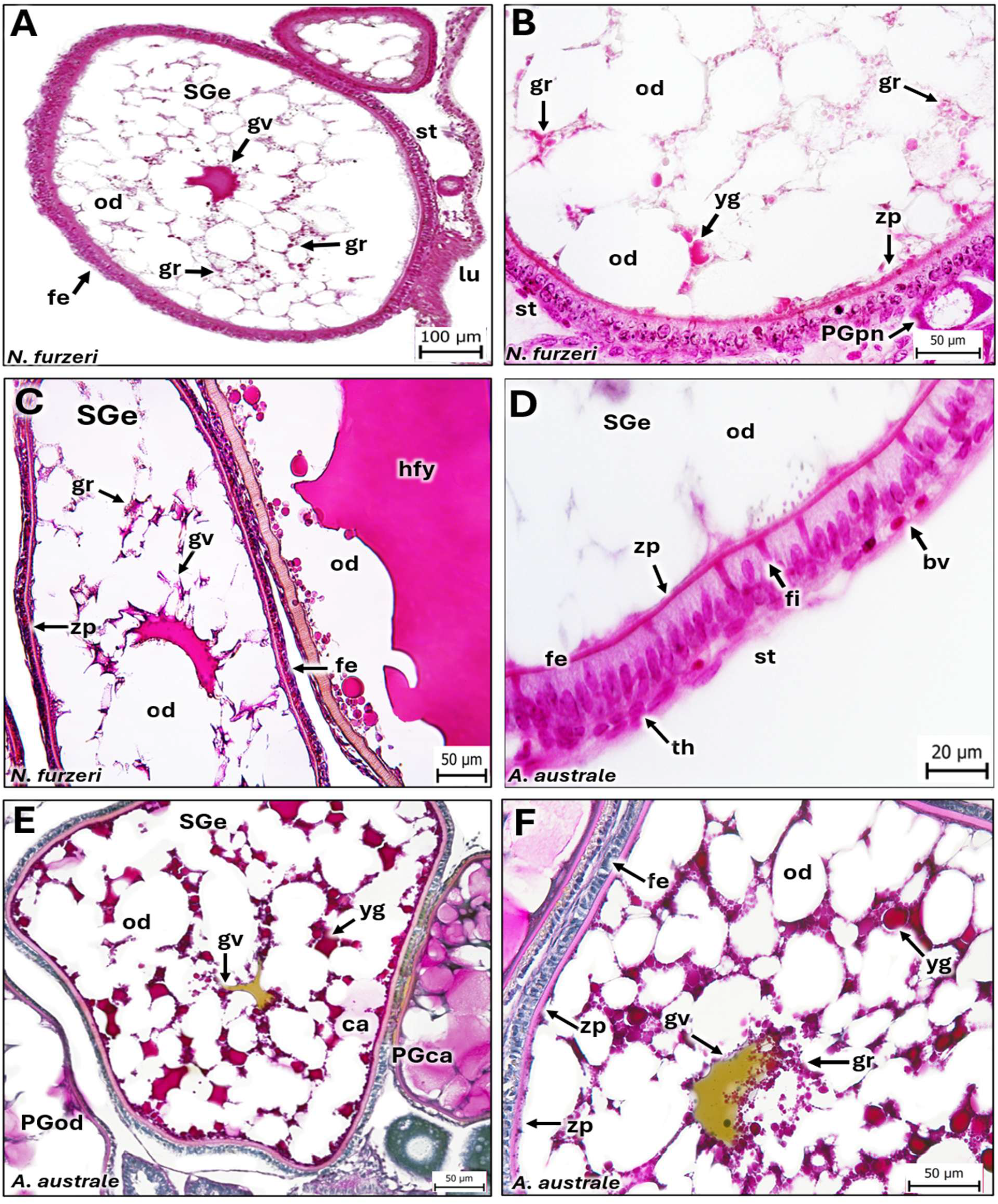
Early Secondary Growth in annual and non-annual killifishes. Early vitellogenesis of the annual killifish *Nothobranchius furzeri* (**A**-**D**) and the non-annual killifish *Aphyosemion australe* (**E**, **F**). (**A**, **E**) Low-magnification views of early Secondary-Growth oocytes (SGe). (**A**-**C**). Small eosinophilic yolk granules first appear throughout the ooplasm between the oil droplets. As the vitellogenesis progresses, these granules progressively fuse into larger yolk globules, resulting in the gradual accumulation of yolk. (**D**) The follicular epithelium differentiates into a tall columnar epithelium, whereas the theca becomes evident as a thin cellular layer associated with connective tissue and blood vessels. The zona pellucida appears as a thin radially striated eosinophilic layer bearing fine projections that extend toward the follicular epithelium. (**E**, **F**) Progressive accumulation and coalescence of yolk granules into yolk globules, together with enlargement of the oil globules, results in displacement of the germinal vesicle toward the oocyte periphery. **Abbreviations:** blood vessel (bv), early Secondary-growth oocyte (SGe), filamentous projections of the zona pellucida (fi), follicular epithelium (fe), germinal vesicle (gv), oil droplet (od), oil globule (og), ovarian lumen (lu), ovarian stroma (st), Primary-Growth cortical alveoli oocyte (PGca), Primary-Growth oil droplet oocyte (PGod), Primary-Growth perinucleolar oocyte (PGpn), theca (th), yolk globule (yg), yolk granule (gr), and zona pellucida (zp), connective tissue (ct). (A-D) Hematoxylin and eosin staining. (E, F) Periodic acid-Schiff and metanil yellow staining.

Similarly, in *A. australe*, small yolk granules and vesicles were initially distributed throughout the ooplasm (Fig. 6A, B, E, F), including the perinuclear region, before progressively coalescing into larger yolk globules in both species (Fig. 6E, F). As a result of this coalescence, the GV became further displaced toward the animal pole. During this stage, the follicular epithelium of *N. furzeri* became columnar, with follicular cells displaying basally positioned nuclei and eosinophilic apical cytoplasm (Fig. 6D). In contrast, the follicular epithelium of *A. australe* remained predominantly cuboidal. Quantitative analysis confirmed that the follicular epithelium was significantly taller *in N. furzeri* than in *A. australe* (Fig. 14F). At this step, the mean height of the follicular epithelium was 10.48 ± 2.13 µm in *N. furzeri* and 5.43 ± 0.95 µm in *A. australe*. During the SGe step, in both species, the zona pellucida increased markedly in thickness (Fig. 6D, E, F). The zona pellucida had a mean thickness of 2.87 ± 1.48 µm in *N. furzeri* and 3.96 ± 1.12 µm in *A. australe*. In *N. furzeri*, projections arising from the zona pellucida extended into the follicular epithelium (Fig. 6D), whereas these filamentous projections were not readily observable in paraffin sections of *A. australe*. The theca remained as a thin layer of flattened cells associated with connective tissue fibers and scattered capillaries (Fig. 6D).

#### 3.4.2. Late secondary growth (SGl) *step*

In *N. furzeri* and *A. australe*, oocytes measured 494.2 ± 63.59 µm and 409.9 ± 55.26 µm, respectively. Both species exhibited a similar pattern of yolk, lipid, and cortical alveoli reorganization at the SGl step. The fusion of yolk globules proceeded from the center toward the periphery of the oocyte. At this step, centrally located yolk globules had already coalesced into a homogeneous fluid yolk mass (Fig. 7A, D, F) or were in process (Fig. 7E), while smaller eosinophilic yolk granules and globules remained in the peripheral ooplasm and were still undergoing progressive coalescence (Fig. 7A, B, F). This process displaced oil droplets and cortical alveoli toward the oocyte periphery and was accompanied by further migration of the GV toward the animal pole (Fig. 7E). The follicular epithelium was columnar in both species at this stage (Fig. 7C, E), with an everage heigh of 9.84 ± 2.03 µm in *N. furzeri* and 8.22 ± 0.76 in *A. australe*. In the latter species, follicular cells exhibited a vesicular appearance in H&E- and toluidine blue-stained sections, with a vesicular basal cytoplasm and a more granular apical region. The theca remained a single layer of flattened cells associated with connective tissue. In both species, the zona pellucida increased considerably in thickness and developed prominent radial striations (Fig. 7B, C, F). The zona pellucida had a mean thickness of 7.20 ± 2.16 µm in *N. furzeri* and 9.86 ± 1.76 µm in *A. australe*. Filamentous projections extending from the zona pellucida were embedded within the follicular epithelium of both species and became particularly conspicuous in *N. furzeri* at this stage, especially in Mallory trichrome-stained sections, in which the filaments stained red (Fig. 7B).

**FIGURE 7.**
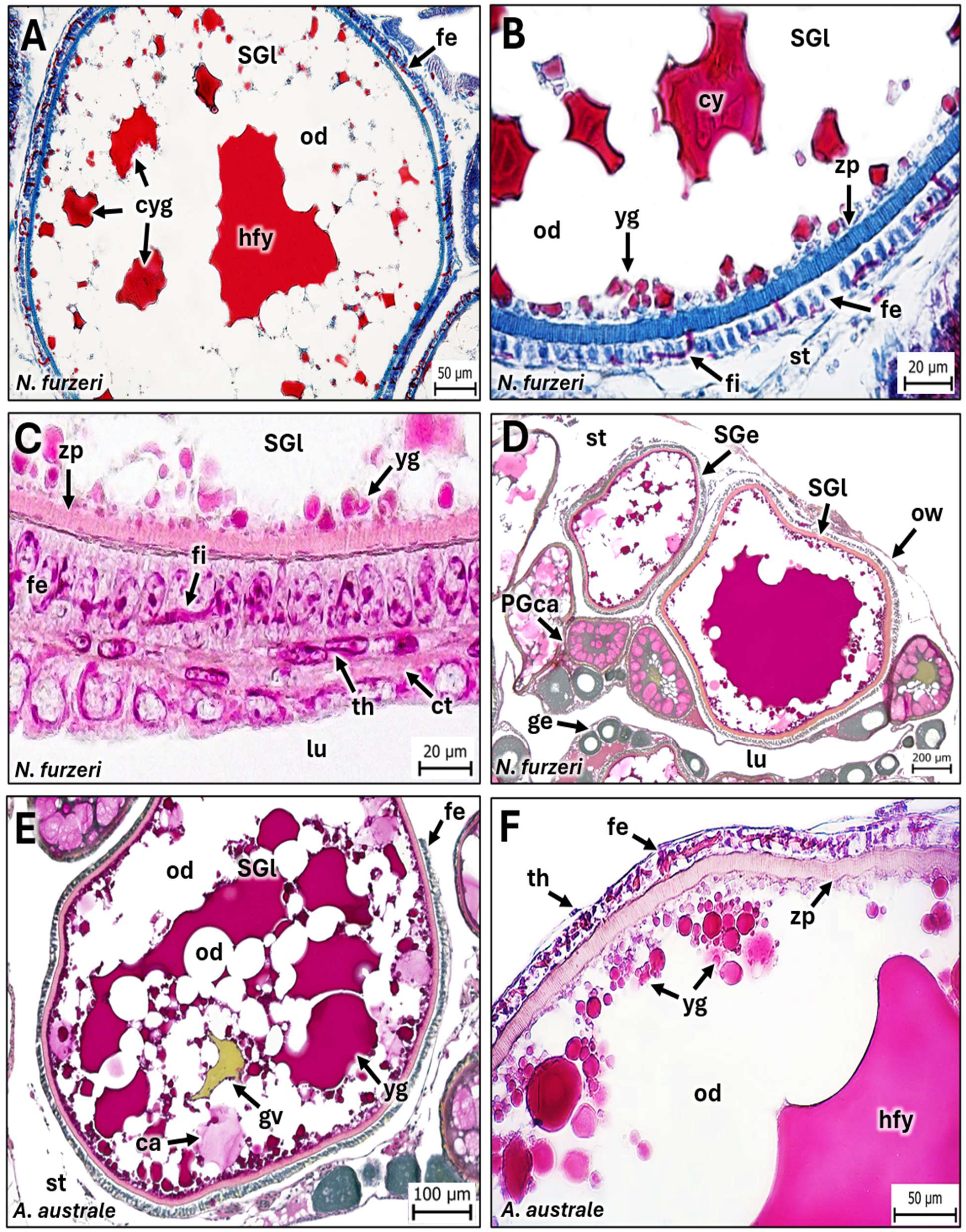
Late Secondary Growth in annual and non-annual killifishes. Mid-vitellogenic oocytes of the annual killifish *Nothobranchius furzeri* (**A**–**C**) and the non-annual killifish *Aphyosemion australe* (**D**–**F**). (**A**, **D**) During the late secondary-growth stage (SGl), the yolk granules deposited during early secondary growth progressively fuse into larger yolk globules, which subsequently coalesce into masses of homogeneous fluid yolk of variable size. Fusion begins in the central ooplasm, resulting in the formation of a large central yolk mass. As yolk accumulation progresses, the expanding yolk masses displace oil globules, cortical alveoli, and germinal vesicle toward the oocyte periphery. Oil globules also fuse into larger droplets. (**C**, **F**) The follicular epithelium remains columnar, whereas the theca persists as a thin layer of flattened cells associated with connective tissue. During this stage, the zona pellucida increases in thickness and exhibits conspicuous radial striations (zona radiata). (**B**, **C**) Fine projections arising from the zona pellucida extend toward the follicular epithelium and are particularly well illustrated in *N. furzeri*. (**D**) Low-magnification view of the ovary showing primary-growth cortical alveoli (PGca), early secondary-growth (SGe), and late secondary-growth (SGl) oocytes. **Abbreviations:** cortical alveoli (ca), coalescent yolk globules (cyg), filamentous projections of the zona pellucida (fi), follicular epithelium (fe), germinal epithelium (ge), germinal vesicle (gv), homogeneous fluid yolk (hfy), late secondary-growth oocyte (SGl), oil globule (og), ovarian lumen (lu), ovarian stroma (st), primary-growth cortical alveoli oocyte (PGca), early secondary-growth oocyte (SGe), theca (th), ovarian wall (ow), yolk globule (yg), and zona pellucida (zp). A, B) Mallory Trichrome. (C, F) Hematoxylin-eosin. (D, E). Periodic acid-Schiff and metanil yellow staining.

#### 3.4.3. Secondary Growth full grown (SGfg)*step*

In *N. furzeri* and *A. australe*, SGfg oocytes reached an average diameter of 689.3 ± 71.86 µm and 809.8 ± 75.39 µm, respectively. In both species, yolk globules coalesced into a homogeneous fluid yolk mass that occupied most of the ooplasm and was surrounded by a peripheral layer of lipids (Figs. 8A–F; 9A–F). A few coalescent yolk globules were observed at the periphery of SGfg oocytes, immersed in the peripheral oil ring (Figs. 8C, D; 9B–D). Cortical alveoli were occasionally observed at the oocyte periphery and were distinguishable only with PAS-trichrome staining (Fig. 9C). The GV was positioned at the oocyte periphery, defining the animal pole (Fig. 8E). In both species, the follicular epithelium consisted of columnar cells with oval nuclei oriented toward the basement membrane (Figs. 8E, F; 9E, F). The height of this layer had a mean height of 10.03 ± 2.33 µm in *N. furzeri* and 12.11 ± 1.78 µm in *A. australe*. The theca was well developed and contained cells with either spherical or flattened nuclei (Figs. 8E, F; 9E, F). The zona pellucida increased considerably in thickness, while retaining its characteristic radial striation. The zona pellucida had a mean thickness of 15.09 ± 2.51 µm in *N. furzeri* and 12.48 ± 1.22 µm in A. australe. Both species exhibited filaments protruding from the zona pellucida into the follicular epithelium; however, these filaments were more conspicuous in *N. furzeri* (Fig. 8C, D) than in *A. australe* (Fig. 9F).

**FIGURE 8.**
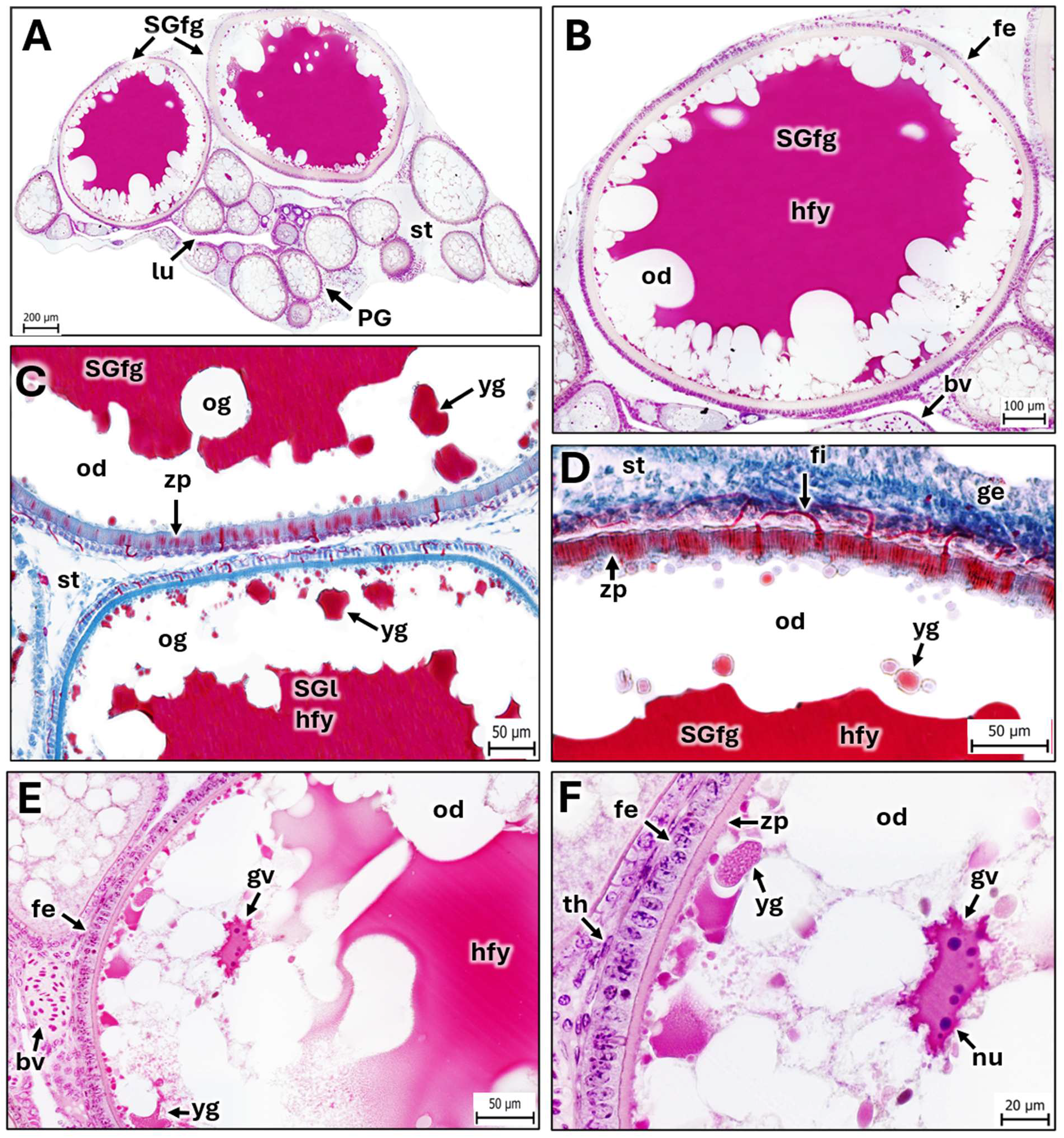
Full-grown Secondary Growth in the annual killifish *Nothobranchius furzeri*. (**A**) Transverse section of an ovary during advanced vitellogenesis showing SGfg oocytes together with primary-growth (PG) and late secondary-growth (SGl) oocytes. (**B**–**D**) At this stage, oocytes reach their maximum size before final maturation, and the ooplasm is almost entirely occupied by a large central mass of homogeneous fluid yolk. Oil globules are displaced toward the oocyte periphery or have fused into a continuous peripheral oil ring. (**F**). The follicular epithelium remains columnar, whereas the theca persists as a thin layer of flattened cells associated with connective tissue. (**C**, **D**) The zona pellucida increases further in thickness while exhibiting prominent radial striations. (**D**) Fine projections arising from the zona pellucida extend toward the follicular epithelium. (**E**, **F**) The germinal vesicle is eccentrically positioned near the oocyte periphery and contains multiple conspicuous nucleoli. **Abbreviations:** blood vessel (bv), follicular epithelium (fe), filamentous projections of the zona pellucida (fi), full-grown secondary-growth oocyte (SGfg), germinal epithelium (ge), germinal vesicle (gv), homogeneous fluid yolk (hfy), late secondary-growth oocyte (SGl), nucleolus (nu), oil globule (og), ovarian lumen (lu), ovarian stroma (st), primary-growth oocyte (PG), theca (th), yolk globule (yg), and zona pellucida (zp). (A, B, E, F) Hematoxylin and eosin staining. (C, D) Mallory’s trichrome staining.

**FIGURE 9.**
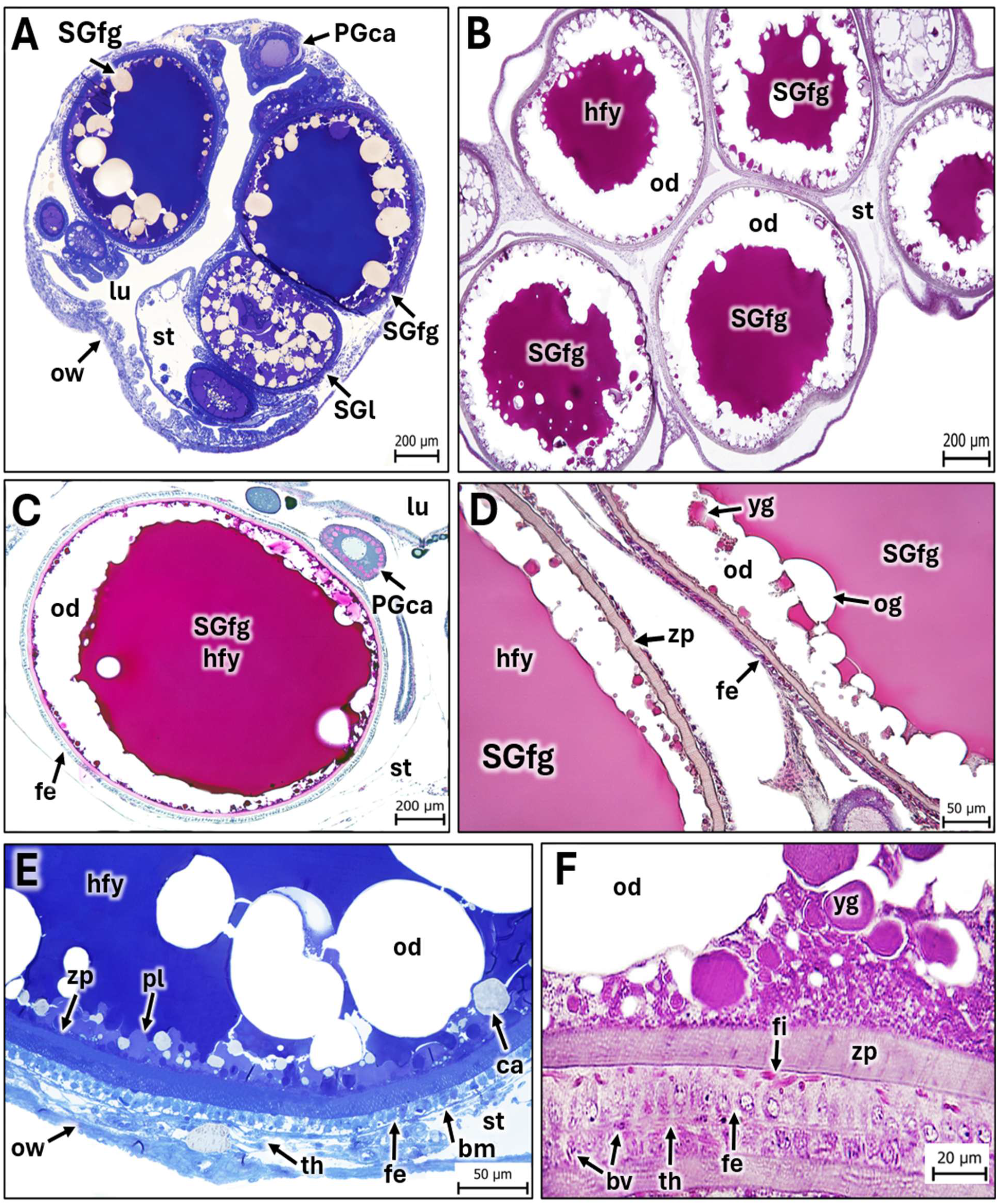
Full-grown Secondary Growth in the non-annual killifish *Aphyosemion australe*. (**A-D**). Representative oocytes at the full-grown Secondary-Growth stage (SGfg). At this stage, yolk accumulation is complete, with the ooplasm largely occupied by a central mass of homogeneous fluid yolk. Oil globules are located at the oocyte periphery or have fused to form a continuous peripheral oil ring. (**E**). The follicular epithelium retains its columnar morphology, with basally positioned nuclei. The zona pellucida increases further in thickness while maintaining its characteristic radial striations. A higher magnification of the peripheral ooplasm reveals residual cortical alveoli, lipid inclusions, and small areas of cytoplasm. (**F**). Fine projections arising from the zona pellucida extend toward the follicular epithelium. The theca remains as a thin layer of flattened cells associated with connective tissue and small blood vessels. **Abbreviations:** blood vessel (bv), filamentous projections of the zona pellucida (fi), follicular epithelium (fe), basement membrane (bm), homogeneous fluid yolk (hfy), oil globule (og), ooplasm (pl), ovarian lumen (lu), ovarian stroma (st), ovarian wall (ow), primary-growth cortical alveoli oocyte (PGca), full-grown secondary-growth oocyte (SGfg), theca (th), yolk globule (yg), and zona pellucida (zp), ovarian wall (ow). (A, E) Semithin sections stained with toluidine blue. (B, D, F) Hematoxylin and eosin staining. (C) Periodic acid–Schiff and metanil yellow staining.

### 3.5. Maturation stage

In *N. furzeri*, mature oocytes had a mean diameter of 868.4 ± 96.04 µm in diameter, whereas in *A. australe* the mean diameter was 1058 ± 93.54 µm. In both species, mature oocytes were readily distinguishable by their irregular shape (Figs. 10A–D; 11A, B) and, in most cases, were either detached from the follicular epithelium immediately prior to ovulation or already ovulated and free within the ovarian cavity (Figs. 10B–D; 11A, B). The zona pellucida had a mean thickness of 21.96 ± 3.16 µm in *N. furzeri* and 12.14 ± 1.43 µm in *A. australe*. The follicular epithelium became an irregular layer of cells (Figs. 10C; 11C, D). Although irregular in nature, in these oocytes, the follicular epithelium had a mean height of 8.96 ± 1.30 µm in *N. furzeri* and 8.84 ± 1.08 µm in *A. australe*. At this stage, the projections of the zona pellucida, which had previously been embedded within the follicular epithelium attained its maximum thickness in *N. furzeri* (Fig. 10E) and *A. australe* (Fig. 11D). In *N. furzeri*, detachment of the oocyte from the follicular epithelium clearly exposed the long filamentous of the zona pellucida, whereas the shorter projections were less conspicuous (Fig. 10E). In *A. australe*, chorionic projections were also observed in paraffin sections, although the long projections were frequently lost during histological processing, leaving predominantly the short, thick projections visible (Figs. 11C, D).

**FIGURE 10.**
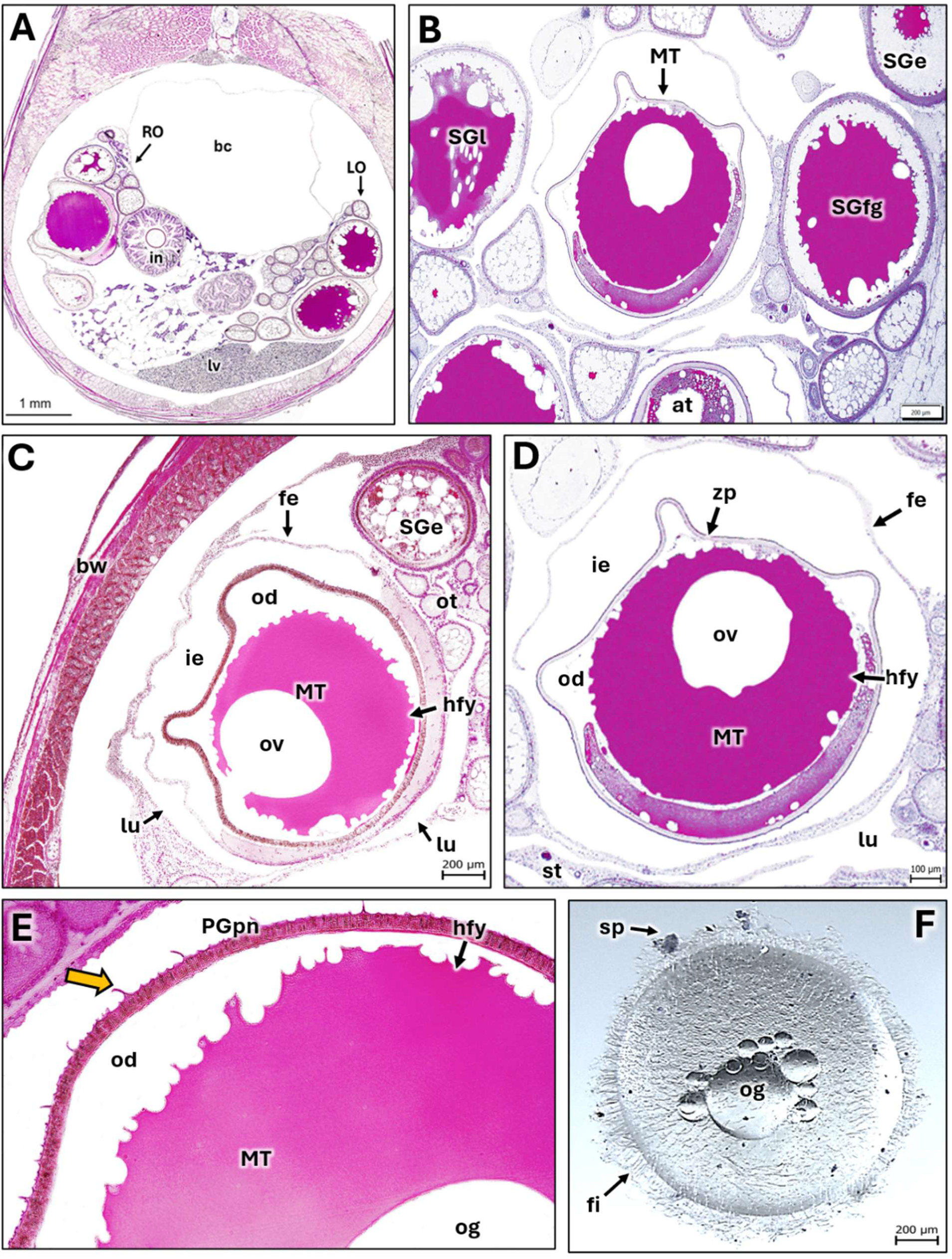
Oocyte maturation in the annual killifish *Nothobranchius furzeri*. Representative stages of oocyte maturation in *N. furzeri*. (**A**) Transverse section of the ovary showing mature oocytes together with secondary-growth oocytes at different stages of development. (**B**) Mature oocyte (MT) surrounded by late secondary-growth (SGl), full-grown secondary-growth (SGfg), and early secondary-growth (SGe) oocytes. (**D**–**E**) During maturation, the ooplasm becomes almost completely occupied by homogeneous fluid yolk, whereas oil globules coalesce into a peripheral oil ring surrounding one or more large central oil globules. The mature oocyte becomes detached from the germinal epithelium and is released into the ovarian lumen. The follicular epithelium remains associated with the oocyte and encloses a conspicuous intrafollicular space. Fine projections of the zona pellucida remain evident (**E**, yellow arrow). F, Visualization of an ovulated unfertilized oocyte of *N. furzeri.* Note the filamentous projections of the chorion that attach well to sand particles. **Abbreviations:** a, atretic oocyte; bc, body cavity; bw, body wall; fe, follicular epithelium; hfy, homogeneous fluid yolk; ie, intrafollicular space; in, intestine; lo, left ovary; lu, ovarian lumen; lv, liver; MT, mature oocyte; og, oil globule; ot, ovarian tissue; RO, right ovary; SGe, early secondary-growth oocyte; SGfg, full-grown secondary-growth oocyte; SGl, late secondary-growth oocyte; st, ovarian stroma; zp, zona pellucida; sp, sand particle; fl, filamentous projections of the chorion; og, oil globules. (A–F) Hematoxylin and eosin staining.

**FIGURE 11.**
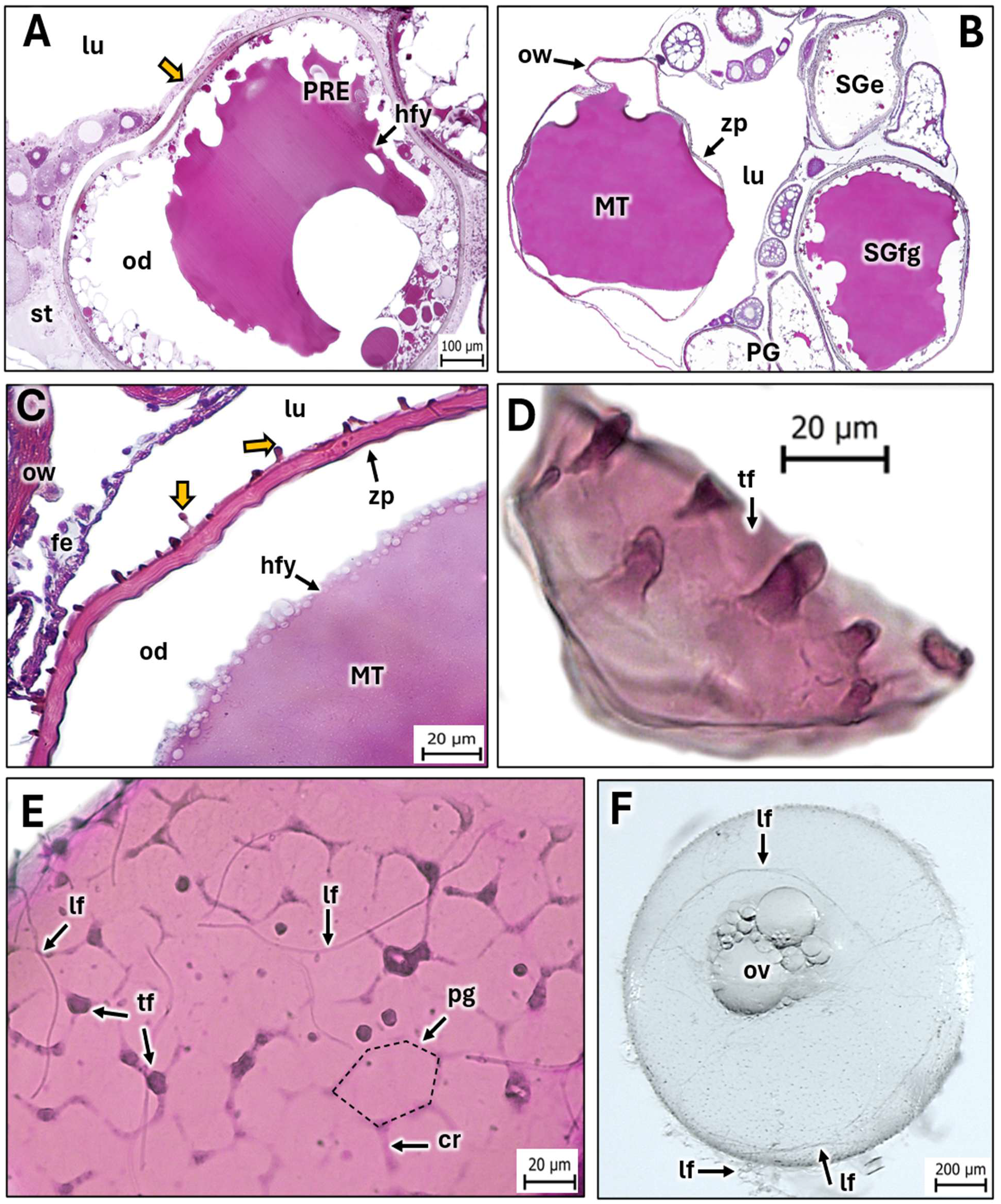
Oocyte maturation and chorion ornamentation in the non-annual killifish *Aphyosemion australe*. Representative mature oocytes and chorion morphology in *A. australe*. (**A**–**C**) Mature oocytes (MT) detached from the follicular epithelium and located within the ovarian lumen. The ooplasm is occupied almost entirely by homogeneous fluid yolk, with oil globules displaced toward the periphery or surrounding one or more large central oil globules. The germinal vesicle is no longer visible. In non-ovulated mature oocytes, the follicular epithelium remains associated with the mature oocyte, enclosing a conspicuous intrafollicular space (A). In ovulated eggs the oocytes are observed free in the ovarian lumen but the remaining follicular epithelium can be seen (C). (**D**–**F**) Surface morphology of the mature chorion. Histological sections parallel to the egg surface reveal polygonal chorionic plates bearing two distinct types of projections: short, thick projections arising from the vertices of the polygonal plates and long, thin filamentous projections emerging from the apex of the short projections. (**E**) Tangential section through the chorion illustrating the polygonal arrangement of the chorionic plates and the distribution of the filamentous projections.(F). An ovulated unfertilized oocyte of *A. australe* showing its long chorionic filaments filaments forming threads. **Abbreviations:** chorionic ridge (cr), follicular epithelium (fe), homogeneous fluid yolk (hfy), long filament (lf), mature oocyte (MT), oil globule (og), ovarian lumen (lu), ovarian stroma (st), ovarian wall (ow), polygonal chorionic plate (pg), primary-growth oocyte (PG), preovulatory follicle (PRE), early secondary-growth oocyte (SGe), full-grown secondary-growth oocyte (SGfg), thick filament (tf), and zona pellucida (zp). (A–H) Hematoxylin and eosin staining.

Occasionally, tangential sections provided a clear view of the external surface of the zona pellucida, revealing the polygonal arrangement of the chorionic plates and the distribution of two distinct types of filamentous projections, long and short (Figs. 11F, G). Once oocyte reaches maturation oogenesis is concluded and it is only followed by ovulation. In Figure 12, we have compared the development of the zona pellucida in *N. furzeri* and *A. australe* throughout SG, when this structure undergoes its most pronounced morphological changes. Finally, in Figure 13, we have summarized the main conserved and divergent events occurring during oocyte development in *N. furzeri* and *A. australe*.

**FIGURE 12.**
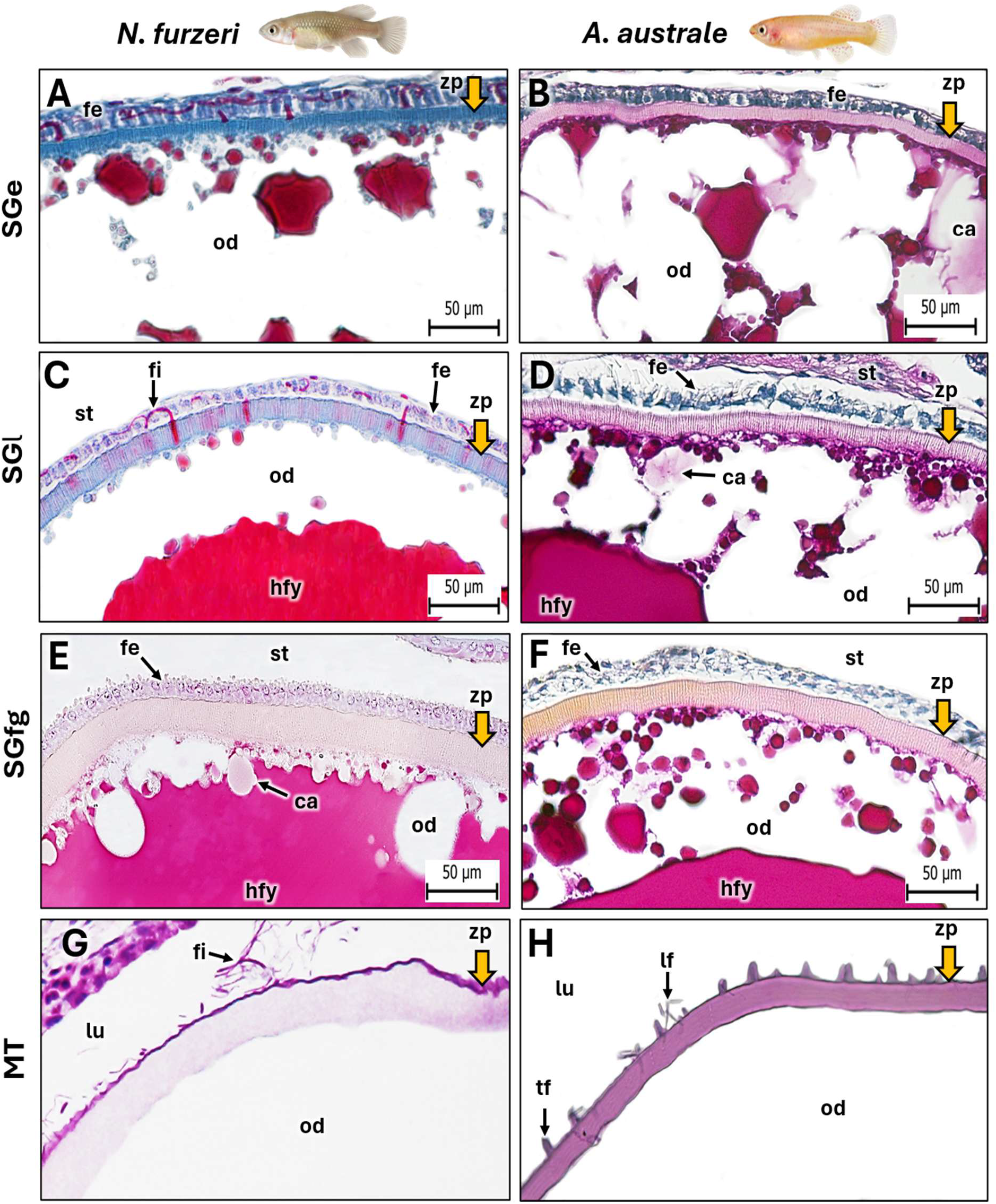
Comparative development of the zona pellucida and chorion in the annual killifish *Nothobranchius furzeri* and the non-annual killifish *Aphyosemion australe*. Changes in the egg envelope throughout oocyte development in *N. furzeri* (**A, C, E, G**) and *A. australe* (**B, D, F, H**). In both species, the zona pellucida during early Secondary Growth (SGe) consisted of a thin radially striated eosinophilic layer. Throughout vitellogenesis, the zona pellucida progressively increases in thickness while maintaining its radial organization and develops fine projections extending toward the follicular epithelium. At the full-grown Secondary-Growth stage (SGfg), the zona pellucida reaches its maximum thickness. During oocyte maturation (MT), the egg envelope becomes multilaminar, giving rise to mature chorion. Throughout oogenesis, the zona pellucida is consistently thicker in *N. furzeri* than in *A. australe*. **Abbreviations:** cortical alveoli (ca), follicular epithelium (fe), homogeneous fluid yolk (hfy), long filament (lf), ovarian lumen (lu), ovarian stroma (st), oil globule (og), oil ring (or), short projection (tf), zona pellucida (zp), and zona pellucida projections (fi). (A, C) Mallory’s trichrome staining. (B, D, F) Periodic acid– Schiff and methanil yellow staining. (E, G, H) Hematoxylin and eosin staining.

**FIGURE 13.**
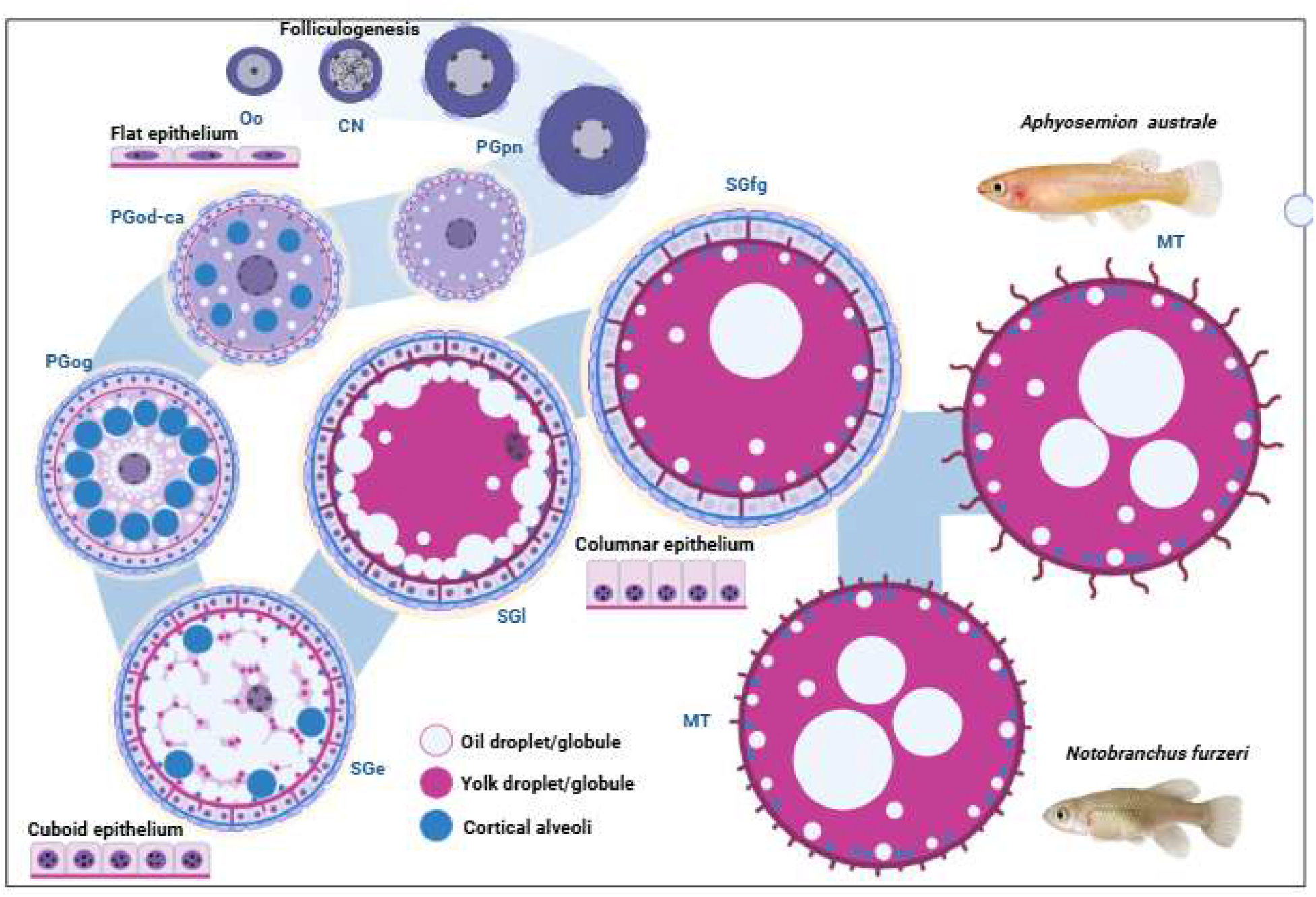
Schematic representation of oogenesis *in Nothobranchius furzeri* and *Aphyosemion australe*. Oocyte development progresses from Oogonia (Oo) and the Chromatin-Nucleolus (CN) and Perinucleolar (PGpn) steps through Primary Growth with cortical alveoli (PGca) and oil globules (PGod), followed by early (SGe), late (SGl), and full-grown Secondary Growth (SGfg) and Maturation (MT). During PG, cortical alveoli and oil droplets progressively accumulate in the ooplasm. Secondary Growth is characterized by yolk deposition and progressive coalescence into homogeneous fluid yolk, accompanied by the redistribution of oil globules and cortical alveoli. Folliculogenesis proceeds concurrently with oocyte growth, with the follicular epithelium transitioning from flattened to cuboidal and subsequently columnar cells. During maturation, the two species exhibit distinct patterns of chorionic ornamentation. Blue, white, and magenta structures represent cortical alveoli, oil droplets/globules, and yolk droplets/globules, respectively. Illustration created in BioRender by Torres-Martínez, A.

### 3.6. Morphometric analysis of oogenesis

Comparison of oocyte diameters between *N. furzeri* and *A. australe* revealed no significant differences at any stage of oogenesis (Fig. 14). In contrast, significant interspecific differences were observed in the height of the follicular epithelium, with follicular cells being taller in *N. furzeri* during the PGod step and during SGe (Fig. 14). The thickness of the zona pellucida also differed significantly between species, being greater in *N. furzeri* (Fig. 14). This difference first became apparent during SGe and persisted through maturation.

**FIGURE 14.**
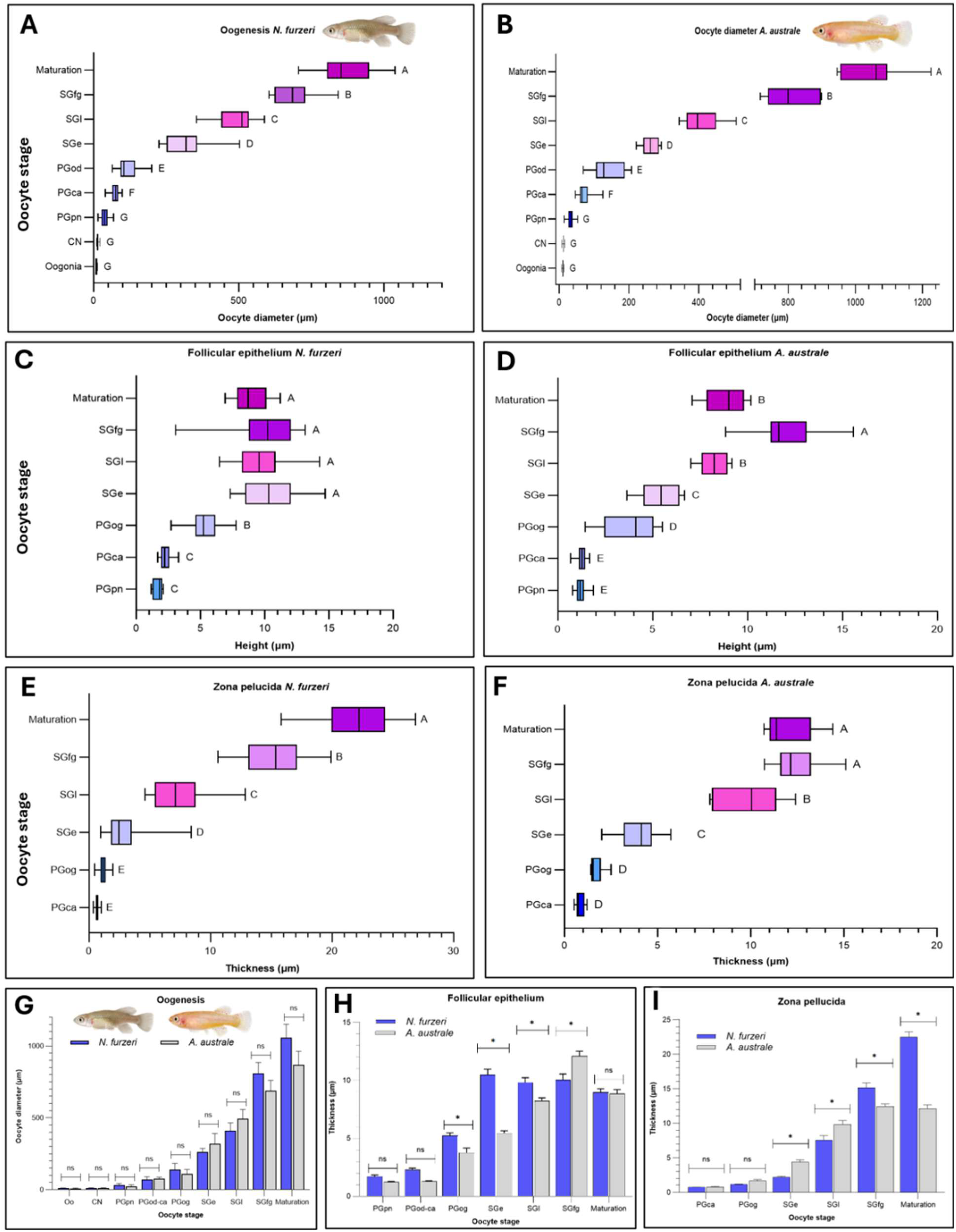
Morphometric comparison of oogenesis in annual and non-annual killifishes. (A–F) Changes in oocyte diameter (A, B), follicular epithelium height (C, D), and zona pellucida thickness (E, F) throughout oogenesis in *Nothobranchius furzeri* (A, C, E) and *Aphyosemion australe* (B, D, F). Different letters indicate statistically significant differences among oocyte developmental stages within each species (P < 0.05). (G–I) Interspecific comparisons of oocyte diameter (G), follicular epithelium height (H), and zona pellucida thickness (I) at corresponding developmental stages. Asterisks indicate statistically significant differences between species at the corresponding oocyte stage (P < 0.05), whereas “ns” indicates no statistically significant difference. Data are presented as boxplots (A–F) and means ± standard error (SE) (G–I). Abbreviations: Oogonia (Oo), Chromatin Nucleolus oocyte (CN), Primary Growth perinucleolar oocyte (PGpn), Primary Growth cortical alveoli oocyte (PGca), Primary Growth oil droplet oocyte (PGod), early Secondary Growth oocyte (SGe), late Secondary Growth oocyte (SGl), Secondary Growth full grown oocyte (SGfg), Mature oocyte (Maturation).

## 4.0. DISCUSSION

### 4.1. Oogonial proliferation

In *Nothobranchius furzeri*, oogonial proliferation was most active during ovarian differentiation, whereas proliferating oogonia were only sparsely distributed throughout the germinal epithelium of adult ovaries. As lifespan progressed, the number of proliferating oogonia declined further and they became rare within the germinal epithelium. Notably, *N. furzeri* follows a repeated spawning strategy throughout adult life, as reported for other annual fishes exhibiting batch spawning, continuous oocyte recruitment, and indeterminate fecundity. This reproductive pattern depends on germ-cell renewal initiated by oogonial proliferation, which is expected to be most active during the regenerating phase (Domínguez-Castanedo et al., 2019). In *Aphyosemion australe*, by contrast, oogonial proliferation was observed at all reproductive ages, although it was considerably more frequent in juvenile individuals, indicating that germ-cell renewal remains active throughout adulthood despite declining with age, as occurs in other fish (Brown-Peterson et al., 2011; Uribe et al., 2014; Wildner et al., 2026).

### 4.2. Morphological changes in the ooplasm before and during Primary Growth (PG, or Previtellogenesis)

It is notable that, despite repeated screening, oocytes corresponding to the Primary Growth One-nucleolus (PGon) and Primary Growth multiple-nucleoli (PGmn) steps were not observed in either species. Nucleolar multiplication begins early during PG, and teleost oocytes commonly develop multiple nucleoli as growth proceeds. During the PGmn step, these nucleoli are initially dispersed throughout the germinal vesicle (GV) and subsequently migrate toward and become associated with the nuclear envelope, resulting in their characteristic peripheral distribution (Wallace, 1981; Thiry & Poncin, 2005; Uribe et al., 2019). The fact that the earliest oocytes identified in *N. furzeri* and *A. australe* already displayed well-developed peripheral nucleoli suggests that nuclear reorganization occurs rapidly during early oogenesis in these killifishes. Indeed, the PGmn step is generally considered transient in fishes and is therefore only rarely observed histologically, likely reflecting the rapid progression of oocytes through this phase (Grier et al., 2009).

Consequently, histologically defined early developmental steps may be short-lived and difficult to capture as discrete categories (Wallace & Selman, 1981; Selman, 1989). This interpretation may be particularly relevant to *N. furzeri*, whose exceptionally rapid life history (Blažek et al., 2013; Vrtílek et al., 2018) could be associated with abbreviated early phases of oocyte development and rapid nucleolar reorganization (Li et al., 2026). Nevertheless, the peripheral localization of nucleoli may not simply reflect rapid developmental progression but could also have functional significance related to their close association with the nuclear envelope (Bir et al., 2026; Li et al., 2026;). The nucleolus is the primary site of rRNA synthesis and ribosomal subunit assembly (Zhang et al., 2018; Kresoja-Rakic and Santoro, 2019), and during oocyte growth, nucleolar activity supports the extensive ribosome biogenesis and accumulation of maternal components required for subsequent development (Li et al., 2026; Zhang et al., 2018) more. Maternal RNAs and ribosomes accumulate during oogenesis in accordance with developmental demands, and the maternal ribosomes produced during this period support protein synthesis during early embryogenesis, before activation of the zygotic genome (Harbey et al., 2013; Shah et al., 2024; Bir et al., 2026).

PG in both species was characterized by marked cytoplasmic basophilia, which is widely associated with increased RNA, ribosome, and protein accumulation (Wallace & Selman, 1981; Uribe & Grier, 2011; Abascal & Medina, 2005). The small vesicular structures observed in the peripheral ooplasm during the PGmn stage are therefore reasonably interpreted as Balbiani body-related aggregates as has been described in other species (Uribe et al., 2012), although histochemical staining alone does not establish their identity unequivocally (Kaufman et al., 2018; Jamieson-Lucy et al., 2022). Oil droplets and oil globules are a major component of fish oogenesis, but their onset and distribution vary among species (Qu et al., 2021; Domínguez-Castanedo, 2024). Their principal role appears to be the storage of neutral lipids, especially triacylglycerol-rich reserves, that support embryonic and larval development; in some taxa they also contribute to buoyancy and prolonged survival under stressful environmental conditions (Cárdenas et al., 2007; Domínguez-Castanedo et al., 2018; Nande et al., 2024). The cortical alveoli on the other hand, are secretory vesicles that release their glycoprotein contents at fertilization to block polyspermy, elevate the chorion, and harden the egg membrane (Bazzoli & Godinho, 1994; Carnevali et al., 2017). The sequence of cortical alveoli and oil-droplet appearance varies among teleosts, and available comparative evidence supports more than one developmental pattern rather than a single conserved order (Bazzoli& Godinho, 1994; Abascal & Medina, 2005; Grier et al., 2009; Uribe et al., 2020). In some species, lipid accumulation begins before cortical alveoli become the dominant cytoplasmic inclusion (Anderson, 1968; Grier et al., 2009; Uribe et al., 2020), as described for Atlantic bluefin tuna *Thunnus thynnus*, in which SG phase starts with a lipid stage before vitellogenesis (Abascal & Medina, 2005) In other species, cortical alveoli appear early, before the onset of oocyte enlargement, as reported *for Syngnathus fuscus* and the killifish *Fundulus heteroclitus* (Anderson, 1968) In a third pattern, both inclusions are deposited simultaneously at the mid and late PG (Uribe et al., 2016; Dominguez-Castanedo, 2024). The latter pattern is consistent with our observations in *N. furzeri* and *A. australe*. In both species, oil droplets begin to accumulate during PG, following the extensive accumulation of cortical alveoli. They initially appear around the GV and progressively increase in abundance until they dominate much of the ooplasm. With the onset and progression of vitellogenesis, the oil droplets are displaced toward the oocyte periphery, where they form a peripheral ring. This sequence indicates that lipid accumulation and redistribution constitute an extended process during oocyte development rather than a late, discrete event (Domínguez-Castanedo, 2018; Domínguez-Castanedo et al., 2019). This pattern is also broadly consistent with descriptions in other atherinomorph and cyprinodontiform fishes, although the relative timing of cortical alveolus formation and lipid accumulation varies among taxa (Parenti & Grier, 2004; Uribe et al., 2016). When comparing *N. furzeri* and *A. australe* with other cyprinodontiform species, considerable interspecific variation is evident in the relative timing of cortical alveolus and oil droplet deposition. In *C. artifrons* and *G. pulchra*, PGca and PGod were not distinguished as separate developmental steps (Domínguez-Castanedo, 2024). In contrast, in the annual Mexican killifish *Millerichthys robustus*, cortical alveolus deposition precedes oil droplet accumulation (Domínguez-Castanedo & Uribe, 2009; Domínguez-Castanedo et al., 2016).In *M. robustus*, cortical alveoli were described as appearing early during oogenesis and are interpreted, together with the zona pellucida and oil droplets, as part of the ovarian assembly of structures associated with embryo survival during diapause (Domínguez-Castanedo, 2016; Domínguez-Castanedo, 2018). Because annualism appears to have originated repeatedly among aplocheiloid killifishes, further comparative work is needed to test whether these cortical-alveolus traits reflect shared ancestry or convergent adaptation to seasonally desiccating habitats (Furness et al., 2015; Furnes, 2016; Thompson et a., 2024). However, caution is warranted when linking early cortical-alveolus formation directly to annualism, because current phylogenetic studies address the repeated evolution of annual life history and diapause rather than the evolution of specific oocyte traits (Furness et al., 2015; Furness, 2016; Thompson et al., 2024). The strongest supported evolutionary inference is that annualism, together with diapause, originated multiple times independently in African and South American killifish lineages (Furness et al., 2015; Thompson et al., 2024) This repeated origin makes convergence plausible, but available evidence is still insufficient to determine whether variation in cortical-alveolus timing reflects shared ancestry, egg physiology, or adaptation to seasonal habitats (Furness, 2016; Thompson et al., 2024). A major obstacle to a stronger evolutionary interpretation is the limited comparative literature on oocyte development and associated egg structures in annual fishes, including killifishes and pupfishes.

### 4.3. Morphological changes in the ooplasm during Secondary Growth (Vitellogenesis)

Secondary Growth (SG) in teleost oocytes is characterized by vitellogenesis, during which yolk is deposited in the ooplasm together with other components required for the nutrition and development of the future embryo (Grier et al., 2009; Lubzens et al., 2010; Uribe et al., 2019). In our target species *N. furzeri* and *A. australe*, SG was characterized by the formation of homogeneous fluid yolk. The pattern closely resembles that described in other Atherinomorpha (Parenti & Grier, 2004; Parenti, 2005), in which individual yolk granules progressively fuse into larger yolk globules that eventually coalesce into a continuous fluid yolk mass occupying most of the ooplasm (Uribe et al., 2016; Torres-Martínez et al., 2017). As vitellogenesis proceeds, the centripetal accumulation and fusion of yolk displace both oil droplets and cortical alveoli toward the periphery of the oocyte (Lubzens et al., 2010).

Although SG is primarily classified according to yolk deposition dynamics, other processes initiated during PG continue throughout this phase, including the accumulation of oil droplets and cortical alveoli (Selman and Wallace, 1989; Uribe et al., 2012). and accordingly, we observed peripheral distribution of the oil droplets and cortical alveoli in *N. furzeri* and *A. australe* from the SGl. Nevertheless, important lineage-specific modifications have evolved within Atherinomorpha. For example, in the Goodeidae *Empetrichthys latos* accumulates only very small oil droplets during SGl and Maturation (Uribe et al., 2012), whereas in the flagfish *Jordanella floridae* the oil droplets remain concentrated around the GV at the animal pole during SGl step (Uribe et al., 2016). The viviparous least killifish *Heterandria formosa* accumulates relatively little yolk but large amounts of oil, a pattern associated with its high degree of matrotrophy (Uribe & Grier, 2011).

Likewise, in the pupfish *Garmanella pulchra*, cortical alveoli accumulate massively beneath the oolemma during SG when compared to another pupfish *Cyprinodon artifrons*, an adaptation associated with the formation of a larger perivitelline space that buffers the effects of desiccation (Domínguez-Castanedo et al., 2024). If there are differences in yolk biogenesis or the accumulation of other inclusions, these were not obvious at the resolution level used here, but they should be explored by other means, such as transmission electron microscopy (Wallace et al., 1990; Kayaba et al., 2001; Jun et al., 2018).

### 4.4. Development of the follicular wall

In addition to cytoplasmic changes, oogenesis in *N. furzeri* and *A. australe* is accompanied by extensive remodeling of structures surrounding the oocyte or the ovarian wall, which includes the zona pellucida, the follicular epithelium, and the theca. There has been some ambiguity in the nomenclature of fish ovarian follicles. In this manuscript, we define the follicular wall as comprising the zona pellucida, the follicular epithelium, and the theca. This usage is practical for comparative morphology, although some authors restrict the term follicle to the oocyte and its surrounding follicle cells and treat the theca as part of a broader follicle complex (Grier, 2000; Quagio-Grassiotto et al., 2011; Viana et al., 2023). In teleosts, the follicular wall typically comprises a zona radiata or zona pellucida apposed to a follicular epithelium, separated externally by a basal lamina or basement membrane from one or more thecal layers (Grier 2000; Jalabert 2005). As with the ooplasm, the morphology of these layers varies widely among teleosts, including differences in zona radiata thickness, pore-canals and microvilli, follicular-cell shape, and thecal organization, and these differences track adaptations associated with egg envelope formation, spawning environment, maternal provisioning, and embryonic protection (Jalabert, 2005; Viana et al., 2023).

#### 4.4.1. The follicular epithelium during the oogenesis of killifishes

Despite the contrasting life histories of *Nothobranchius furzeri*, which inhabits ephemeral seasonal pools, and *Aphyosemion australe*, which occupies more permanent aquatic habitats, the overall development of the follicular wall appears highly conserved. Annual killifishes repeatedly evolved traits associated with survival in temporary waters, including diapause and desiccation-resistant eggs, yet many aspects of ovarian follicle development remain shared across teleosts and likely reflect strong developmental constraint rather than ecological divergence (Jalabert, 2005; Quagio-Grassiotto et al., 2011; Domínguez-Castanedo et al., 2016; Furness, 2016). In both killifish species studied here, the follicular epithelium was formed by a monolayer of flattened cells during early oogenesis (from CN stage to the PGmn step) and became cuboidal (PGod step) and then columnar as oocyte growth progressed (from SGe to SGfg steps)., Similar transitions from flattened to cuboidal and finally columnar epithelial cells have been described in *Cyprinodon artifrons* and *Garmanella pulchra* (Domínguez-Castanedo & Uribe, 2019; Domínguez-Castanedo et al., 2024), as well as in *Millerichthys robustus* (Domínguez-Castanedo & Uribe, 2019). A broadly comparable pattern has also been reported for *Melanorivulus rossoi*, although the available material did not clearly document the SG steps (Cassel et al., 2023). As a matter of fact, it matches a broadly conserved teleost pattern during PG (previtellogenesis) and SG (vitellogenesis) (Patiño & Sullivan, 2002; Ravaglia & Maggesse, 2003; Uribe et al., 2016). These morphological changes coincide with the intense accumulation of cytoplasmic reserves during the late PG and SG (Wallace & Selman, 1981; 1990), consistent with the proposed role of follicular cells in mediating the transport of nutrients and other materials to the growing oocyte. This epithelial differentiation coincided with the phase of rapid oocyte enlargement and reserve accumulation, consistent with the established role of follicular tissues in supporting vitellogenic growth. In teleosts, vitellogenesis depends on hepatic vitellogenin synthesis, receptor-mediated uptake by the oocyte, and coordinated follicular endocrine activity, while lipids from non-vitellogenin sources can also contribute substantially to follicle growth (Patiño & Sullivan, 2002; Jalabert, 2005;). Ultrastructural studies further show that material reaches the oocyte through stromal, thecal, granulosa, and zona radiata compartments before being internalized by micropinocytosis, supporting the interpretation that changes in epithelial height track changing transport demands during oocyte growth (Matsuyama et al., 1991; Grove & Wourms, 1994). Although the developmental sequence was shared, the follicular epithelium was consistently taller in *N. furzeri* than in *A. australe* from the oil-globule step through the SGl step, and this difference disappeared after SG. The functional significance of this transient difference remains unresolved, but comparative work suggests that variation in follicular-cell morphology can accompany differences in reproductive tactics, egg traits, and oocyte physiology, including changes in zona radiata thickness, thecal organization, and follicular-layer thickness among teleost lineages (Viana et al., 2023). In annual killifishes, follicular specializations linked to harsh seasonal environments are more evident in structures such as the zona pellucida, cortical alveoli, and secondary envelope than in the basic sequence of epithelial differentiation (Domínguez-Castanedo et al., 2016).

#### 4.4.2. Thickness of the zona pellucida and its adaptive functions in killifish from contrasting environments

Among the components of the follicular wall, the zona pellucida is one of the most morphologically variable structures in killifishes and teleosts more broadly, varying in thickness, stratification, and external ornamentation across species (Thompson et al., 2017; Litscher & Wassarman, 2018; Messaddeq et al., 2018). Despite this broader diversity, its developmental sequence was similar in *N. furzeri* and *A. australe*. In both species, the zona pellucida developed from a thin layer during early oogenesis, thickened during SG, and acquired a more complex multilaminar organization during oocyte maturation, consistent with the general teleost pattern in which the egg envelope first appears as an oocyte-derived layer and progressively becomes stratified during growth (Tesoriero, 1977; Ravaglia & Maggesse, 2003). This interpretation is also consistent with comparative studies showing that zona pellucida proteins are synthesized during oogenesis and assembled into a specialized extracellular matrix that supports egg-envelope formation and remodeling, beyond providing physical protection to ovulated eggs (Patiño & Sullivan, 2002; Sano et al., 2022). Quantitative analysis nevertheless showed that the zona pellucida was significantly thicker in *N. furzeri* than in *A. australe*. That difference is biologically plausible, because comparative studies across teleosts show substantial interspecific variation in zona radiata thickness and indicate that this layer can covary with reproductive tactics, egg traits, and spawning environment (Viana et al., 2023). For example, in viviparous fishes the thickness of the zona pellucida has been reduced at the minimum to allow internal fertilization (Riehl & Greven, 1993) or maternal supply to the embryos during intrafollicular development (Uribe & Grier, 2011), while annual fishes are known to have thicker zona pellucida (Domínguez-Castanedo & Uribe, 2019; 2024). At the same time, the presence of a relatively thick egg envelope in both species is notable given their contrasting life histories. *N. furzeri* is an annual killifish from ephemeral pools whose embryos survive seasonal desiccation in diapause, whereas non-annual killifishes retain the ancestral longer-lived life history associated with more permanent aquatic habitats. A thicker zona pellucida in *N. furzeri* is therefore consistent with adaptation to annualism, but it does not by itself show that this trait is unique to annual species or specifically caused by selection for desiccation tolerance. One possibility is that a relatively thick egg envelope represents an ancestral condition retained within Nothobranchiidae, with further modification of its thickness in lineages such as *Nothobranchius* following adaptation to ephemeral environments. However, this hypothesis cannot be distinguished from independent or convergent evolution based on the two species examined here. That caution is reinforced by the comparative killifish literature. Across aplocheiloid killifishes, chorionic characters show strong homoplasy and repeated structural convergence among lineages (Thompson et al., 2017). Chorionic ornamentation has been extensively documented using light microscopy (Arezo et al., 2005) and scanning electron microscopy (Loureiro & De Sa, 1996; Thompson et al., 2017), yet no simple ecological or phylogenetic pattern has emerged. A comparative study of seasonal and non-seasonal cyprinodontiforms likewise found species-level variation in chorion thickness and ornamentation, but no obvious phylogenetic or ecological relationship between chorion structure and seasonality in the sampled taxa (Messaddeq et al., 2018). Even at the molecular level, teleost zona pellucida genes show lineage-specific evolution while retaining core functions related to egg-envelope hardening and hatching (Sano et al., 2022). Together, these findings make retained ancestry, ecological modification, and convergent evolution, all plausible explanations for the thicker zona pellucida in *N. furzeri*. The annual-killifish literature supports a similarly restrained interpretation. Annualism evolved independently in multiple killifish lineages, and parallel evolution under similar selective pressures is a recurring pattern in these fishes (Furness, 2016; Tozzini & Cellerino, 2020; Polačik et al., 2023). Moreover, extreme egg-envelope specializations are documented in at least some annual taxa. In *Millerichthys robustus*, for example, the zona pellucida is exceptionally thick and is associated with additional envelope elaborations interpreted as structures important for embryo survival during prolonged diapause (Domínguez-Castanedo et al., 2016). However, this does not establish that all thick envelopes in nothobranchiids are homologous or that they all arose for the same ecological reason. A more plausible interpretation is that a relatively thick egg envelope may represent a shared nothobranchiid tendency that has been further modified in some annual lineages, including *Nothobranchius*, after adaptation to ephemeral environments.

#### 4.4.3. Adornment of the zona pellucida and its adaptive functions in killifish from contrasting environments

Greater interspecific differences were evident in the external ornamentation of the egg envelope. In *N. furzeri*, the chorion bore elongated hook-like projections interspersed with short hair-like structures, whereas *A. australe* exhibited short mushroom-like projections interspersed with long filaments; in both species, these ornamentations became evident during oogenesis and persisted in ovulated oocytes. This interpretation is consistent with comparative work showing that killifish chorions display substantial variation in external surface adornments, while the internal chorion also varies in thickness and stratification among species (Messaddeq et al., 2017; Thompson et al., 2017; Messaddeq et al., 2018). Comparative analyses across aplocheiloid and other teleost lineages likewise show wide diversity in the shape and arrangement of chorionic projections, including filaments, clustered filament-bearing structures, cruciform hooks, anchor-shaped hooks, and net-like outer sheets, indicating that egg-envelope surface morphology is evolutionarily labile (Mooi, 1990). Similar diversity is also seen outside killifishes, where egg capsules or chorions show species-specific ornamentation patterns that are useful for description and comparison, but not always straightforward to homologize across taxa (Berrada-Rkhami et al., 1990; Merret et al., 1996;

Nieves-Uribe et al., 2020). The marked difference in chorionic ornamentation between *N. furzeri* and *A. australe*, despite their close taxonomic affinity within Nothobranchiini, therefore fits a broader pattern of high chorion diversification in killifishes and other egg-laying taxa (Costa, 2018; Nieves-Uribe et al., 2020) Proposed functions for these structures remain plausible but incompletely tested. Comparative evidence supports roles in adhesion and anchoring, because some fish eggs possess specialized attaching filaments that unite neighboring eggs and anchor them to the female gonoduct, and external traits such as hooks or spines can facilitate substrate adhesion in dormant eggs from temporary-water taxa (Hart et al., 1984; Meyer-Milne et al., 2021). Environmental interaction is also a credible function, since egg-envelope thickness and topographic morphology can reflect physicochemical conditions, and ornamentation categories sometimes correlate with habitat distribution (Berrada-Rkhami & Gabrion, 1990; Merret et al., 1996). However, killifish comparative studies found no obvious phylogenetic or ecological relationship between external adornments and seasonality in the sampled species, and broader work on egg-surface morphology warns that similar structures are often non-homologous and difficult to polarize phylogenetically (Mooi, 1990; Messaddeq et al., 2017; 2018). In annual species such as *N. furzeri*, specialized surface structures therefore remain consistent with functions such as anchoring within substrate or mediating dispersal through adhesion to mud or animal vectors, but those interpretations still require direct experimental testing (Meyer-Milne et al., 2021).

### 4.5. Future directions: Where do we go from here?

Overall, the comparison between *N. furzeri* and *A. australe* indicates that the general sequence of follicular wall formation is conserved, whereas features such as chorion thickness, surface ornamentation, and cortical alveoli dynamics appear more evolutionarily labile and likely track differences in reproductive ecology (Domínguez-Castanedo et al., 2016; Furness, 2016; Thompson et al., 2017). This interpretation fits the broader biology of killifishes, in which annual lineages repeatedly evolved embryo-based persistence in ephemeral habitats, while non-annual relatives occupy more stable aquatic environments, creating a strong comparative framework for testing how oocyte and egg-envelope traits relate to environmental challenge (Furness, 2015; 2016; 2018). Comparative evidence also suggests caution: chorionic characters in aplocheiloid killifishes are highly homoplastic, and similar egg-envelope structures can evolve convergently rather than reflecting simple shared ancestry (Thompson et al., 2017; Polačik et al., 2023). A morphologically detailed description of oogenesis is therefore valuable for more than descriptive purposes. It establishes homologous oocyte stages for cross-species comparison, which is essential in a group where annualism evolved repeatedly and where developmental trajectories can diverge in response to both shared and lineage-specific selective pressures (Furness, 2015; Polačik et al., 2023). It also provides the anatomical framework needed to investigate how annualism-related structures arise during oogenesis. In annual killifishes, cortical alveoli, zona pellucida, oil droplets, and secondary envelope specializations are all implicated in embryo survival, and their origin can be traced to distinct phases of follicle development (Domínguez-Castanedo et al., 2016; 2018).

Moreover, the staging framework that we presented here is especially useful because developmental responses in annual killifish are regulated by both maternal inputs and environmental cues, including temperature, light, incubation medium, and seasonal conditions (Furness et al., 2015; Domínguez-Castanedo et al., 2018; 2022; Pérez-Atehortúa et al., 2023). Classical interpretations have emphasized mechanical roles in attachment and protection, but studies of egg envelops in other animal groups, especially insects, show that these structures also protect against biotic and abiotic stress, including predation, parasitism, drought, adverse temperatures, and solar radiation (Rezende et al., 2016; Li et al., 2024). In fishes, the chorion likewise functions as more than a passive barrier: it protects the oocyte and embryo, can show bactericidal and fungicidal properties, and its thickness can increase resistance to pathogen invasion (Berois et al., 2011; Songe et al., 2016; Pérez-Atehortúa et al., 2023). The structural diversity of the killifish chorion may therefore reflect a similarly broad range of functions, although direct functional tests are still limited (Berois et al., 2011; Papa et al., 2019; Pérez-Atehortúa et al., 2023).

Lastly, beyond clarifying the biogenesis of structures associated with annualism, this study provides a framework for identifying specific oocyte stages, which is necessary for investigating the molecular and physiological regulation of oocyte development in killifishes (Berois et al., 2011). This staging is important because fish chorion formation depends on regulated synthesis of choriogenins, which can occur in the liver, the oocyte, or both, and because chorion quality is shaped by environmental and nutritional factors that affect embryo survival (Pérez-Atehortúa et al., 2023). In annual killifishes specifically, few studies have linked egg-envelope structure to its molecular basis; one example is the identification of two hormone-regulated egg-envelope proteins in *Austrolebias charrua*, which helps clarify the regulation and function of this unusually resistant envelope (Papa et al., 2019). More generally, there remain substantial gaps in understanding the molecular and physiological mechanisms that connect intracytoplasmic structures such as cortical alveoli with the formation of thicker, stress-resistant chorions (Berois et al., 2011; Pérez-Atehortúa et al., 2023).

## CONFLIC OF INTEREST

The authors declare no conflicts of interest.

## ACKNOWLEGMENTS

We would like to thank Michaela Vazačová and Zuzana Havlíková-Linhartová for their care of the animals throughout this study.

## AUTHOR CONTRIBUTIONS

**Aarón Torres-Martínez:** Conceptualization, Formal analysis, Investigation, Visualization, Writing – Original Draft Preparation. Essaikiammal Sodalai Muthu Konar: Writing – Review & Editing**. Martin Pšenička**: Funding adquisition, Resources, Writing – Review & Editing. **Roman Franěk:** Conceptualization, Investigation, Funding acquisition, Resources, Visualization.

## ACKNOWLEDGMENTS

The authors have nothing to report.

## FUNDING INFORMATION

This research was co-funded by Czech Science Foundation (22-01781O), Ministry of Education, Youth and Sports of the Czech Republic projects Biodiversity (CZ.02.1.01/0.0/0.0/16_025/0007370) and OP JAK Aquaculture for Future (CZ.02.01.01/00/23_021/0012616).

## Notes

### Competing Interest Statement

The authors have declared no competing interest.

### Summary of Updates

Figures 4 and 5 inadvertently showed the same image. We have replaced Figure 5 with the correct image and adjusted the placement of the figures within the manuscript. We apologize for the inconvenience and would appreciate it if you could replace the current PDF with the revised version.

## REFERENCES

Abascal, F. & Medina, A. (2005) Ultrastructure of oogenesis in the bluefin tuna, *Thunnus thynnus*. Journal of Morphology, 264, 149–160. 10.1002/jmor.10325

Anderson, E. (1968) Cortical alveoli formation and vitellogenesis during oocyte differentiation in the pipefish, *Syngnathus fuscus*, and the killifish, *Fundulus heteroclitus*. Journal of Morphology, 125(1), 23–59. 10.1002/jmor.1051250103

Arezo, M.J., Pereiro, L. & Berois, N. (2005) Early development in the annual fish *Cynolebias viarius*. Journal of Fish Biology, 66, 1357–1370. 10.1111/j.0022-1112.2005.00688.x

Bazzoli, N. & Godinho, H.P. (1994) Cortical alveoli in oocytes of freshwater Neotropical teleost fish. Italian Journal of Zoology, 61, 301–308. 10.1080/11250009409355899

Bankhead, P., Loughrey, M.B., Fernández, J.A., Dombrowski, Y., McArt, D.G., Dunne, P.D. et al. (2017) QuPath: open source software for digital pathology image analysis. Scientific Reports, 7, 16878. 10.1038/s41598-017-17204-5

Berois, N., Arezo, M. & Papa, N. (2011) Gamete interactions in teleost fish: the egg envelope. Basic studies and perspectives as environmental biomonitor. Biological Research, 44, 119–124. 10.4067/S0716-97602011000200002

Berrada-Rkhami, O. & Gabrion, C. (1990) The fine structure of the embryonic envelopes before and after hatching in bothriocephalids: physiological and ecological significance. Parasitology Research, 76, 251–262. 10.1007/BF00930822

Bir, J., De Cerio, O.D., Ortiz-Zarragoitia, M. & Cancio, I. (2026) Differential regulation of ribogenesis genes throughout oogenesis in thicklip grey mullets, *Chelon labrosus*, preparing the maternal contribution to embryo development. PLOS ONE, 21, e0356468. 10.1371/journal.pone.0356468

Borisov, V., Shkil, F.N., Seleznev, D. & Smirnov, S. (2023) Is African non-annual killifish *Fundulopanchax gardneri* (Teleostei; Cyprinodontiformes; Nothobranchiidae) true non-annual? Developmental Dynamics, 253, 490–507.10.1002/dvdy.668

Brown-Peterson, N., Wyanski, D.M., Saborido-Rey, F., Macewicz, B. & Lowerre-Barbieri, S. (2011) A standardized terminology for describing reproductive development in fishes. Marine and Coastal Fisheries, 3, 52–70. 10.1080/19425120.2011.555724

Campbell, B., Dickey, J., Beckman, B., Young, G., Pierce, A., Fukada, H. et al. (2006) Previtellogenic oocyte growth in salmon: relationships among body growth, plasma insulin-like growth factor-1, estradiol-17β, follicle-stimulating hormone and expression of ovarian genes for insulin-like growth factors, steroidogenic-acute regulatory protein and receptors for gonadotropins, growth hormone, and somatolactin. Biology of Reproduction, 75, 34–44. 10.1095/biolreprod.105.049494

Cao, Y.Q., Wang, Y., Zhao, Y., Zhang, J., He, X., Xie, P. et al. (2023) Transfer of the *zp3a* gene results in changes in egg adhesiveness and buoyancy in transgenic zebrafish. Zoological Research, 44, 259– 268. 10.24272/j.issn.2095-8137.2022.330

Cárdenas, R., Chávez, M., González, J.L., Aley, P., Espinosa, J. & Jiménez-García, L. (2007) Oocyte structure and ultrastructure in the Mexican silverside fish *Chirostoma humboldtianum* (Atheriniformes: Atherinopsidae). Revista de Biología Tropical, 56, 1825–1835. 10.15517/rbt.v56i3.5715

Carnevali, O., Candelma, M., Sagrati, A., Pignalosa, P., Giorgini, E. & Gioacchini, G. (2019) Macromolecular characterization of swordfish oocytes by FTIR imaging spectroscopy. Scientific Reports, 9, 1–9. 10.1038/s41598-019-45065-7

Cassel, M., Severo-Neto, F., Mehanna, M. Ferreira, A. (2023) Morphological and temporal variations in the gametogenic development of the killifish *Melanorivulus rossoi* (Cyprinodontiformes, Rivulidae) in an area under crop influence. Zoomorphology, 142, 447–452. 10.1007/s00435-023-00615-6

Costa, W.J.E.M. & Leal, F. (2009) Egg surface in the Neotropical seasonal killifish genus *Leptolebias* (Teleostei: Aplocheiloidei: Rivulidae). Vertebrate Zoology, 59, 25–29. 10.3897/vz.59.e30942

Domínguez-Castanedo, O., Uribe, M.C. & Rosales-Torres, A.M. (2016) Morphological development of the structures related to annualism in the ovarian follicle of the killifish *Millerichthys robustus* (Costa, 1995) (Teleostei: Cyprinodontiformes). Journal of Morphology, 277, 1219–1230. 10.1002/jmor.20570

Domínguez-Castanedo, O., Valdesalici, S. & Rosales-Torres, A. (2018) Reproductive biology of annual killifishes and its relationship with embryonic survival during diapause: *Millerichthys robustus* (Cyprinodontiformes: Cynolebiidae) as an integrative model. Journal of Applied Ichthyology, 34, 793–800. 10.1111/jai.13623

Domínguez-Castanedo, O. & Uribe, M.C. (2019) Ovarian structure, folliculogenesis and oogenesis of the annual killifish *Millerichthys robustus* (Cyprinodontiformes: Cynolebiidae). Journal of Morphology, 280, 316–328.10.1002/jmor.20945

Domínguez-Castanedo, O. (2024) Ovarian structure and oogenesis of two endemic pupfishes from the Yucatán Peninsula: morphofunctional traits related with an estuarine life history. Zoomorphology, 143, 443–458. 10.1007/s00435-024-00653-8

Franěk, R., Šindelka, R., Torres-Martínez, A., Štundl, J., Konar, E.S.M., Siqueira-Silva, D.E. et al. (2026) A lifespan atlas of the killifish testis defines a pronounced mid-age remodeling shift during testicular aging. BMC Biology, 24, 195. 10.1186/s12915-026-02689-2

Furness, A.I., Reznick, D., Springer, M. & Meredith, R. (2015) Convergent evolution of alternative developmental trajectories associated with diapause in African and South American killifish. Proceedings of the Royal Society B: Biological Sciences, 282, 20142189. 10.1098/rspb.2014.2189

Furness, A.I. (2016) The evolution of an annual life cycle in killifish: adaptation to ephemeral aquatic environments through embryonic diapause. Biological Reviews, 91. 10.1111/brv.12194

Gao, L., Franěk, R., Tichopád, T., Rodina, M., Gela, D., Šindelka, R., et al. (2025) Mitochondria as indispensable yet replaceable components of germ plasm: insights into PGCs specification in sturgeons. Reproduction, 169, e240441. 10.1530/REP-24-0441

Grier, H. (2000) Ovarian germinal epithelium and folliculogenesis in the common snook, *Centropomus undecimalis* (Teleostei: Centropomidae). Journal of Morphology, 243, 265–281. 10.1002/%28SICI%2910974687%28200003%29243%3A3%3C265%3A%3AAID-JMOR4%3E3.0.CO%3B2-I

Grier, H.J., Uribe, M.C. & Patiño, R. (2009) The ovary, folliculogenesis, and oogenesis in teleosts. In: Jamieson, B.G.M. (Ed.) Reproductive biology and phylogeny of fishes. New Hampshire, USA: Science Publishers, pp. 25–84.

Grier, H. (2012) Development of the follicle complex and oocyte staging in red drum, *Sciaenops ocellatus* Linnaeus, 1766 (Perciformes, Sciaenidae). Journal of Morphology, 273, 801–829. 10.1002/jmor.20034

Grier, H., Porak, W., Carroll, J. & Parenti, L. (2018) Oocyte development and staging in the Florida bass, Micropterus floridanus (LeSueur, 1822), with comments on the evolution of pelagic and demersal eggs in bony fishes. Copeia, 106, 329–345. 10.1643/cg-17-679

Grove, B. & Wourms, J. (1994) Follicular placenta of the viviparous fish, Heterandria formosa: II. Ultrastructure and development of the follicular epithelium. Journal of Morphology, 220, 167–184. 10.1002/jmor.1052200206

Harvey, S.A., Sealy, I., Kettleborough, R.N.W., Fényes, F., White, R.J., Stemple, D. et al. (2013) Identification of the zebrafish maternal and paternal transcriptomes. Development, 140, 2703–2710. 10.1242/dev.095091

Hart, N., Pietri, R. & Donovan, M. (1984) The structure of the chorion and associated surface filaments in *Oryzias*—evidence for the presence of extracellular tubules. Journal of Experimental Zoology, 230, 273–296. 10.1002/jez.1402300213

Hernández-Gómez, R., Contreras-Sánchez, W.M., Hernández-Franyutti, A.A., Perera-García, M.A. & Torres-Martínez, A. (2019). Reproductive cycle of Aplodinotus grunniens females (Rafinesque, 1819) in the Usumacinta River, Mexico. Latin. American. Journal of Aquatic Research, .47, 612-625. 10.3856/vol47-issue4-fulltext-4

Hilker, M., Salem, H. & Fatouros, N.E. (2023) Adaptive plasticity of insect eggs in response to environmental change. Annual Review of Entomology, 68, 451–469. 10.1146/annurev-ento-120120-100746

Honorato-Sampaio, K., Prado, P.S., Sato, Y., Bazzoli, N. & Rizzo, E. (2015) Comparative morphology of the oocyte surface and early development in four characiformes from the São Francisco River, Brazil. Journal of Morphology, 276, 1258–1272. 10.1002/jmor.20416

Jalabert, B. (2005) Particularities of reproduction and oogenesis in teleost fish compared to mammals. *Reproduction*, Nutrition and Development, 45, 261–279. 10.1051/rnd:2005019

Jun, J.-C., Gang, H.W. & Lee, K.-Y. (2018) Ultrastructural studies on oocyte differentiation and vitellogenesis in the oocytes of female *Kareius bicoloratus* in western Korea. Development & Reproduction, 22, 213–223. 10.12717/dr.2018.22.3.213

Kayaba, T., Takeda, N., Adachi, S. & Yamauchi, K. (2001) Ultrastructure of the oocytes of the Japanese eel *Anguilla japonica* during artificially induced sexual maturation. Fisheries Science, 67, 870–879. 10.1046/j.1444-2906.2001.00335.x

Kresoja-Rakic, J. & Santoro, R. (2019) Nucleolus and rRNA gene chromatin in early embryo development. Trends in Genetics, 35, 868–879. 10.1016/j.tig.2019.06.005

Leow, C. & Piller, K.R. (2025) Life history-dependent brain transcriptomic signatures in nothobranchids: insights into aging, neurogenesis, and life history evolution. Integrative Organismal Biology, 7, 1–14. 10.1093/iob/obaf016

Li, R., McKown, G., Tsuchiya, D., Mattingly, M., Galligos, A.M., Diez, M. et al. (2026) Nucleolar dynamics during oogenesis. bioRxiv [Preprint]. 10.64898/2026.05.19.726235

Li, T.-H., Wang, X., Desneux, N., Wang, S. & Zang, L. (2024) Egg coverings in insects: ecological adaptation to abiotic and biotic selective pressures. Biological Reviews, 100, 99–112. 10.1111/brv.13130

Litscher, E.S. & Wassarman, P.M. (2018) The fish egg’s zona pellucida. Current Topics in Developmental Biology, 130, 275–305. 10.1016/bs.ctdb.2018.01.002

Loureiro, M. & de Sá, R.O. (1996) External morphology of the chorion of the annual fishes *Cynolebias* (Cyprinodontiformes: Rivulidae). Copeia, 1996(4), 1016–1022. 10.2307/1447669

Lubzens, E., Young, G., Bobe, J. & Cerdà, J. (2010) Oogenesis in teleosts: how fish eggs are formed. General and Comparative Endocrinology, 165, 367–389. 10.1016/j.ygcen.2009.05.022

Lubzens, E., Bobe, J., Young, G. & Sullivan, C. (2017) Maternal investment in fish oocytes and eggs: the molecular cargo and its contributions to fertility and early development. Aquaculture, 472, 107– 143. 10.1016/j.aquaculture.2016.10.029

Martin, K. & Podrabsky, J. (2017) Hit pause: developmental arrest in annual killifishes and their close relatives. Developmental Dynamics, 246, 858–866. 10.1002/dvdy.24507

Matsuyama, M., Nagahama, Y. & Matsuura, S. (1991) Observations on ovarian follicle ultrastructure in the marine teleost, *Pagrus major*, during vitellogenesis and oocyte maturation. Aquaculture, 92, 67– 82. 10.1016/0044-8486(91)90009-V

Merrett, N. & Barnes, S.H. (1996) Preliminary survey of egg envelope morphology in the Macrouridae and the possible implications of its ornamentation. Journal of Fish Biology, 48, 101–119. 10.1111/j.1095-8649.1996.tb01422.x

Messaddeq, N., Hergueux, J., Weickert, J. & Romand, R. (2018) Study of the chorion of seasonal and non-seasonal Africa and Neotropical oviparous Cyprinodontiforme fishes. Environmental Biology of Fishes, 101, 287–299. 10.1007/s10641-017-0698-7

Mokhtar, D.M. (2024) Diversity and dynamics of fish ovaries: insights into reproductive strategies, hormonal regulation, and ovarian development. Histology and Histopathology, 40, 283–295. 10.14670/hh-18-802

Mooi, R.D. (1990) Egg surface morphology of pseudochromoids (Perciformes: Percoidei), with comments on its phylogenetic implications. Copeia, 1990, 455–475. 10.2307/1446351

Nakamura, S., Kobayashi, K., Nishimura, T., Higashijima, S. & Tanaka, M. (2010) Identification of germline stem cells in the ovary of the teleost medaka. Science, 328, 1561–1563. 10.1126/science.1185473

Nande, M., Pérez, M. & Presa, P. (2024) The embryo-oil drop assembly: the timing and morphology of a critical event for fish early-life history survival. Scientific Reports, 14, 1–14. Available from: 10.1038/s41598-024-57429-9

Nieves-Uribe, S., Flores-Gallardo, A. & Llorente-Bousquets, J. (2020) Chorion exploration in the tribe Anthocharidini (Lepidoptera: Pieridae) and their possible importance in its systematics. Zootaxa, 4868, 151–207. 10.11646/zootaxa.4868.2.1

Papa, N.G., Chalar, C., Berois, N. et al. (2019) Annual killifish: an approach to the choriogenins of *Austrolebias charrua* egg envelope. Environmental Biology of Fishes, 102, 829–844. 10.1007/s10641-019-00861-y

Parenti, L.R. (2005) The phylogeny of atherinomorphs: evolution of a novel fish reproductive system. In: Grier, H.J. & Uribe, M.C. (Eds.) Viviparous fishes. Homestead, FL: New Life Publications.

Parenti, L.R. & Grier, H.J. (2004) Evolution and phylogeny of gonad morphology in bony fishes. Integrative and Comparative Biology, 44, 333–348. 10.1093/icb/44.5.333

Patiño, R. & Sullivan, C. (2002) Ovarian follicle growth, maturation, and ovulation in teleost fish. Fish Physiology and Biochemistry, 26, 57–70. 10.1023/A:1023311613987

Pérez-Atehortúa, M., Hernández, A.J., Dantagnan, P., Silva, M., Risopatrón, J., Farías, J. et al. (2023) Chorion in fish: synthesis, functions and factors associated with its malformations. Aquaculture Reports, 30, 1–10. 10.1016/j.aqrep.2023.101590

Podrabsky, J., Carpenter, J. & Hand, S.J. (2001) Survival of water stress in annual fish embryos: dehydration avoidance and egg envelope amyloid fibers. *American Journal of Physiology-Regulatory*, Integrative and Comparative Physiology, 280, R123–R131. 10.1152/ajpregu.2001.280.1.R123

Podrabsky, J. & Arezo, M. (2017) Annual killifishes as model systems for advancing understanding of evolution and developmental biology. Developmental Dynamics, 246, 778. 10.1002/dvdy.24594

Polačik, M., García, D., Arezo, M., Papa, N., Schlueb, H., Blanco, D. et al. (2023) Embryonic development of natural annual killifish populations of the genus *Austrolebias*: evolutionary parallelism and the role of environment. Freshwater Biology. A, 68, 1726–1738. 10.1111/fwb.14161

Qu, J., Li, R., Xie, Y., Liu, Y., Liu, J. & Zhang, Q. (2021) Differential transcriptomic profiling provides new insights into oocyte development and lipid droplet formation in Japanese flounder (*Paralichthys olivaceus*). Aquaculture, 550, 1–11. Available from: *Aquaculture*. 10.1016/j.aquaculture.2021.737843

Quagio-Grassiotto, I., Grier, H., Mazzoni, T., Nóbrega, R. & de Arruda Amorim, J.P. (2011) Activity of the ovarian germinal epithelium in the freshwater catfish, *Pimelodus maculatus* (Teleostei: Ostariophysi: Siluriformes): germline cysts, follicle formation and oocyte development. Journal of Morphology, 272, 1290-1306. 10.1002/jmor.10981

Quintero-Hunter, I., Grier, H. & Muscato, M. (1991) Enhancement of histological detail using metanil yellow as counterstain in periodic acid Schiff’s hematoxylin staining of glycol methacrylate tissue sections. Biotechnic & Histochemistry, 66, 169–172. 10.3109/10520299109109964

Ravaglia, M. & Maggese, M. (2003) Ovarian follicle ultrastructure in the teleost *Synbranchus marmoratus* (Bloch, 1795), with special reference to the vitelline envelope development. Tissue & Cell, 35, 9–17. 10.1016/S0040-8166(02)00098-8

Reichard, M. (2016) The evolutionary ecology of African annual fishes. In: Berois, N., García, G. & de Sá, R.O. (Eds.) Annual fishes: life history strategy, diversity, and evolution. Boca Raton, FL: CRC Press.

Rezende, G.L., Vargas, H.C.M., Moussian, B. & Cohen, E. (2016) Composite eggshell matrices: chorionic layers and sub-chorionic cuticular envelopes. pp. 325–366. 10.1007/978-3-319-40740-1_9

Riehl, R. & Greven, H. (1993) Fine structure of egg envelopes in some viviparous goodeid fishes, with comments on the relation of envelope thinness to viviparity. Canadian Journal of Zoology, 71, 91– 97. 10.1139/z93-014

Riehl, R. & Patzner, R. (1998) Minireview: the modes of egg attachment in teleost fishes. Italian Journal of Zoology, 65, 415–420. 10.1080/11250009809386857

Rizzo, E., Sato, Y., Barreto, B.P. & Godinho, H.P. (2002) Adhesiveness and surface patterns of eggs in Neotropical freshwater teleosts. Journal of Fish Biology, 61, 615–632. 10.1111/j.1095-8649.2002.tb00900.x

Sano, K., Kawaguchi, M., Katano, K., Tomita, K., Inokuchi, M., Nagasawa, T. et al. (2017) Comparison of egg envelope thickness in teleosts and its relationship to the sites of ZP protein synthesis. Journal of Experimental Zoology Part B: Molecular and Developmental Evolution, 328, 240–258. 10.1002/jez.b.22729

Sano, K., Shimada, S., Mibu, H., Taguchi, M., Ohsawa, T., Kawaguchi, M. et al. (2022) Lineage-specific evolution of zona pellucida genes in fish. Journal of Experimental Zoology Part B: Molecular and Developmental Evolution, 338, 181–191. 10.1002/jez.b.23122

Shah, A.N., Leesch, F., Lorenzo-Orts, L., Grundmann, L., Novatchkova, M., Haselbach, D. et al. (2024) A dual ribosomal system in the zebrafish soma and germline. bioRxiv [Preprint]. 10.1101/2024.08.29.610041.

Tesoriero, J. (1977) Formation of the chorion (zona pellucida) in the teleost, Oryzias latipes. I. Morphology of early oogenesis. Journal of Ultrastructure Research, 59, 282–291. 10.1016/S0022-5320(77)90040-5

Thiry, M. & Poncin, P. (2005) Morphological changes of the nucleolus during oogenesis in oviparous teleost fish, *Barbus barbus* (L.). Journal of Structural Biology, 152, 1–13. 10.1016/j.jsb.2005.07.006

Thompson, A.W., Black, A., Huang, Y., Shi, Q., Furness, A.I., Braasch, I. et al. (2024) Transcriptomic data supports phylogenetic congruence and reveals genomic changes associated with the repeated evolution of annualism in aplocheiloid killifishes (Cyprinodontiformes). Molecular Phylogenetics and Evolution, 201, 1–14. 10.1016/j.ympev.2024.108209

Thompson, A.W., Furness, A.I., Stone, C., Rade, C. & Ortí, G. (2017) Microanatomical diversification of the zona pellucida in aplocheiloid killifishes. Journal of Fish Biology, 91, 126–143. 10.1111/jfb.13332.o

Tichopád, T., Vetešník, L., Šimková, A., Rodina, M., Franěk, R., Pšenička, M. (2020) Spermatozoa morphology and reproductive potential in F1 hybrids of common carp (*Cyprinus carpio*) and gibel carp (*Carassius gibelio*). Aquaculture, 521, 735092. 10.1016/j.aquaculture.2020.735092

Torres-Martínez, A., Hernández-Franyutti, A., Uribe, M.C. & Contreras-Sánchez, W.M. (2017) Ovarian structure and oogenesis of the extremophile viviparous teleost *Poecilia mexicana* (Poeciliidae) from an active sulfur spring cave in Southern Mexico. Journal of Morphology, 278, 1667–1681. 10.1002/jmor.20740

Torres-Martínez, A., Tichopád, T., Pšenička, M. & Franěk, R. (2026) Triploidy alters hormonal and paracrine signaling to promote male development in zebrafish. Molecular and Cellular Endocrinology, 615, 1–22. 10.1016/j.mce.2026.112740

Tozzini, E.T. & Cellerino, A. (2020) Nothobranchius annual killifishes. EvoDevo, 11. 10.1186/s13227-020-00170-x

Tyler, C. & Sumpter, J. (1996) Oocyte growth and development in teleosts. Reviews in Fish Biology and Fisheries, 6, 287–318. 10.1007/BF00122584

Uribe, M.C., de la Rosa-Cruz, G. & Grier, H.J. (2014) Proliferation of oogonia and folliculogenesis in the viviparous teleost *Ilyodon whitei* (Goodeidae). Journal of Morphology, 275, 1004–1015. 10.1002/jmor.20277

Uribe, M.C. & Grier, H.J. (2011) Oogenesis of microlecithal oocytes in the viviparous teleost *Heterandria formosa*. Journal of Morphology, 272, 241–257.10.1002/jmor.10912

Uribe, M.C., Grier, H.J., García-Alarcón, A. & Parenti, L.R. (2016) Oogenesis: from oogonia to ovulation in the flagfish, *Jordanella floridae* Goode and Bean, 1879 (Teleostei: Cyprinodontidae). Journal of Morphology, 277, 1339–1354. 10.1002/jmor.20580

Uribe, M.C., Grier, H.J. & Hernández-Franyutti, A.A. (2019) Ovaries and eggs. In: Formicki, K. & Kirschbaum, F. (Eds.) The histology of fishes. Boca Raton, FL: CRC Press.

Uribe, M.C., Grier, H.J. & Parenti, L.R. (2012) Ovarian structure and oogenesis of the oviparous goodeids *Crenichthys baileyi* (Gilbert, 1893) and *Empetrichthys latos* Miller, 1948 (Teleostei, Cyprinodontiformes). Journal of Morphology, 273, 371–387.10.1002/jmor.11028

Viana, I.K., Ferreira, M., Mendes, Y., Silva, B.R.M., Gonçalves, L.A.B. & Rocha, R. (2023) Follicular complex may predict reproductive tactics in siluriform fishes. Heliyon, 9, 1–9. 10.1016/j.heliyon.2023.e16204

Vrtílek, M., Žák, J., Pšenička, M. & Reichard, M. (2018) Extremely rapid maturation of a wild African annual fish. Current Biology, 28, R822–R824.10.1016/j.cub.2018.06.031

Wallace, R.A. & Selman, K. (1981) Cellular and dynamic aspects of oocyte growth in teleosts. Integrative and Comparative Biology, 21, 325–343. 10.1093/icb/21.2.325

Wallace, R.A. & Selman, K. (1990) Ultrastructural aspects of oogenesis and oocyte growth in fish and amphibians. Journal of Electron Microscopy Technique, 16, 175–201. 10.1002/jemt.1060160302

Wiegand, M. (1996) Composition, accumulation and utilization of yolk lipids in teleost fish. Reviews in Fish Biology and Fisheries, 6, 259–286. 10.1007/bf00122583

Wildner, D., Grier, H. & Quagio-Grassiotto, I. (2013) Female germ cell renewal during the annual reproductive cycle in Ostariophysian fish. Theriogenology, 79, 709–724. 10.1016/j.theriogenology.2012.11.028

Wourms, J.P. (1976) Annual fish oogenesis. I. Differentiation of the mature oocyte and formation of the primary envelope. Developmental Biology, 50, 338–354. 10.1016/0012-1606%2876%2990156-1

Wourms, J.P. & Sheldon, H. (1976) Annual fish oogenesis. II. Formation of the secondary egg envelope. Developmental Biology, 50, 355–366. 10.1016/0012-1606%2876%2990157-3

Zhao, Y., Xue, G., Peng, Y.-B., Zhang, J., Chen, F., Wang, Y.-K. et al. (2025) Ultrastructure and transcriptomic analysis reveal alternative pathways of zona radiata formation in *Culter alburnus* with different spawning habits. Biology, 14, 987. 10.3390/biology14080987.

